# A Generic Force Field for Simulating Native Protein Structures Using Dissipative Particle Dynamics

**DOI:** 10.1101/2021.08.16.456428

**Authors:** Rakesh Vaiwala, K. Ganapathy Ayappa

## Abstract

A coarse-grained force field for molecular dynamics simulations of native structures of proteins in a dissipative particle dynamics (DPD) framework is developed. The parameters for bonded interactions are derived by mapping the bonds and angles for 20 amino acids onto target distributions obtained from fully atomistic simulations in explicit solvent. A dual-basin potential is introduced for stabilizing backbone angles, to cover a wide spectrum of protein secondary structures. The backbone dihedral potential enables folding of the protein from an unfolded initial state to the folded native structure. The proposed force field is validated by evaluating structural properties of several model peptides and proteins including the SARS-CoV-2 fusion peptide, consisting of *α*-helices, *β*-sheets, loops and turns. Detailed comparisons with fully atomistic simulations are carried out to assess the ability of the proposed force field to stabilize the different secondary structures present in proteins. The compact conformations of the native states were evident from the radius of gyration as well as the high intensity peaks of the root mean square deviation histograms, which were found to lie below 0.4 nm. The Ramachandran-like energy landscape on the phase space of backbone angles (*θ*) and dihedrals (*ϕ*) effectively captured the conformational phase space of *α*-helices at ∼(*ϕ* = 50°, *θ* = 90°) and *β*-strands at ∼(*ϕ* = ±180°, *θ* = 90° − 120°). Furthermore, the residue-residue native contacts are also well reproduced by the proposed DPD model. The applicability of the model to multidomain complexes is assessed using lysozyme as well as a large *α* helical bacterial pore-forming toxin, cytolysin A. Our studies illustrate that the proposed force field is generic, and can potentially be extended for efficient in-silico investigations of membrane bound polypeptides and proteins using DPD simulations.

## 1 Introduction

Molecular dynamics (MD) simulations have been extensively used to uncover the understanding of processes that occur at the molecular scale and has emerged as an indispensable computational tool to study a wide variety of systems with atomistic as well as mesoscopic detail. Although accurate in its description, studying the structure and dynamics of large biomolecular complexes for longer times with atomistic resolution using all-atom simulations is computationally prohibitive. Coarse-grained (CG) approaches, with a reduction in the number of degrees of freedom to enhance conformational sampling, allow assessing processes that occur at physiologically relevant spatial and temporal scales. Consequently, there has been a rapid development of CG models and multiscale simulation methods in last few decades for simulating simple fluids, ionic liquids, biomembranes, proteins and transmembrane protein assemblies. ^1–11^

With an increase in virulent bacterial and viral infections, CG modelling of biological systems involving membrane-protein complexes has attracted considerable attention. ^11–14^ As a result, numerous CG protein models at various levels of reduced resolution ranging from one to five CG beads mapped to represent an amino acid residue have been developed. ^15,16^ Depending on the level of description used to characterize the protein and solvent, models vary in their ability to capture protein folding, conformational dynamics and prediction of the native states. These CG models of proteins are structure-based, ^17–24^ knowledge-based, ^25–30^ and physics-based, ^31–37^ with varying degrees of structural accuracy, model transferability and thermodynamic consistency. The MARTINI, OPEP, PRIMO, UNRES, and SIRAH are among some of the popular protein force fields for CG-MD simulations. ^5,15,16,38–40^

Dissipative particle dynamics (DPD) is a particle based simulation method, which allows one to examine the underlying system at mesoscopic length and time scales. ^41,42^ DPD has been extensively used to study biomembranes, polymers and polyelectrolytes, and it has been shown to predict the scaling behaviour of several systems. ^43–51^ However, DPD has been used to a lesser extent to study systems involving peptides and proteins. The DPD simulation framework is attractive for simulation of proteins, since dynamics are naturally evolved in the canonical ensemble in the presence of explicit solvent particles.

In an early study, Vishnyakov *et al*. ^52^ introduced a novel approach to model secondary structures of polypeptides using DPD simulations. The *α*-helices and *β*-sheet structures were stabilized using dissociable Morse bonds between 1-3 as well as 1-5 amino acid residues in the protein sequence. The stiffness constant for the Morse bonds, the charge on the residues and the hydrophobicity were varied to capture a given secondary structure, without the need for amino acid specific interaction potentials. The authors also point out that the presence of multiple Morse bonds due to their spherical symmetry could cause peptides to adopt unphysical conformations. ^52^ Following the MARTINI scheme for assigning bead types to protein residues, a CG mapping scheme for amino acids has been proposed for simulating antimicrobial peptides using the DPD methodology. ^53,54^ The repulsive force parameters for backbone beads are specific to the secondary structures being simulated, and the Morse potential between pairs of backbone beads has been used to stabilize *α*-helices and *β*-sheet structures.

The knowledge-based protein force field proposed by Peter *et al*. ^55^ is a polarizable model for simulating proteins in the DPD framework. A pair of oppositely charged Drude particles, which are transparent to other CG beads, has been employed to mimic electrostatic polarization of the peptide backbone. Being a polarizable force field, the protein model is computationally expensive in a polarizable solvent model. In a DPD based model developed by Truszkowski *et al*., ^56^ a protein force field has been presented with a CG mapping scheme based on underlying molecular fragments. For instance, the side chains of serine and threonine were represented by the CG beads that correspond to methanol and ethanol fragments, respectively. In absence of anisotropic interactions in the model, the structural rigidity of the protein was maintained by harmonic bonds between the backbone beads, similar to the elastic dynamic network models used in MARTINI simulations, ^57^ adding to the computational overhead. This also precludes simulating protein folding from an unfolded state as well as secondary structure transitions.

In a very recent DPD study on folding of polyalanine into its native structure, the *α*-helix was stabilized by optimizing the repulsive force constant for the alanine residues that form the helical structure using the native contacts derived from atomistic simulations. ^58^ The dihedral potential was used to stabilize the helical structure. Nevertheless, the parameter set is polyalanine specific, and therefore the parameters need to be re-optimized for different proteins with varying secondary structures.

The above studies reveal that although DPD models have been used to model specific protein structures, an efficient and generic protein force field for simulating native protein structures in the DPD formalism is yet to be developed. Additionally, it is desirable to develop a protein force field which can be integrated with existing DPD models for phospholipid membranes. ^43,53^ In an attempt to bridge this gap, we present a complete set of force field parameters for 20 amino acids for simulating polypeptides and proteins using DPD. The native conformations, namely *α*-helix, rigid turn, flexible loop, *β*-hairpin and other extended structures, are preserved by introducing backbone dihedrals. Not restricted to a specific protein or protein structure, the proposed force field for bonded and non-bonded interactions is generic and can be applied to a wide range of peptides and protein structures. Unlike previous force fields developed for DPD simulations, the force field presented here does not require a priori knowledge of the presence of distinct secondary structures such as *α*-helices and *β*-sheets. Additionally the model is computationally inexpensive when compared with the polarizable protein models, ^55^ and the elastic network based models. ^56^ A fact that the emergence of covid-19 shook the world ^64^ has led a rising interest in protein-membrane interactions, and a full trimeric spike protein model for SARS-CoV-2 is also now available with our CG force field for such studies in the DPD framework.

## 2 Simulation methodology

In this section the protocols for simulating the amino acid sequences in atomistic as well as CG frameworks are described. The peptides chosen for parametrizing and validating the DPD force field for proteins are depicted in Fig. 1. These peptides include all the standard 20 amino acids, encompassing distinct structures such as *α*-helices, *β*-sheets, turns and flexible loops. Some of these peptides such as Trpzip (1LE1), Trpcage (1L2Y), 1HDN, 1J4M and lysozyme have been previously used for developing various protein force fields. ^7,55,58,65–68^ The peptides shown in Fig. 1A are used to parametrize the DPD force field, while the peptides and proteins in Fig. 1B are used to test the robustness of the proposed force field. In addition to the smaller peptides we carried out DPD simulations of complex multidomain bacterial toxin cytolysin A (1QOY) and lysozyme (3TXJ) to demonstrate the versatility of the proposed force field (Fig. 1B). The monomer of cytolysin A (ClyA), a 34 kDa protein expressed by *Escherichia coli*, takes part in membrane assisted oligomerization during pore formation. We also modify the force field to describe disulfide bonds between cysteine residues while simulating lysozyme.

**Fig. 1.**
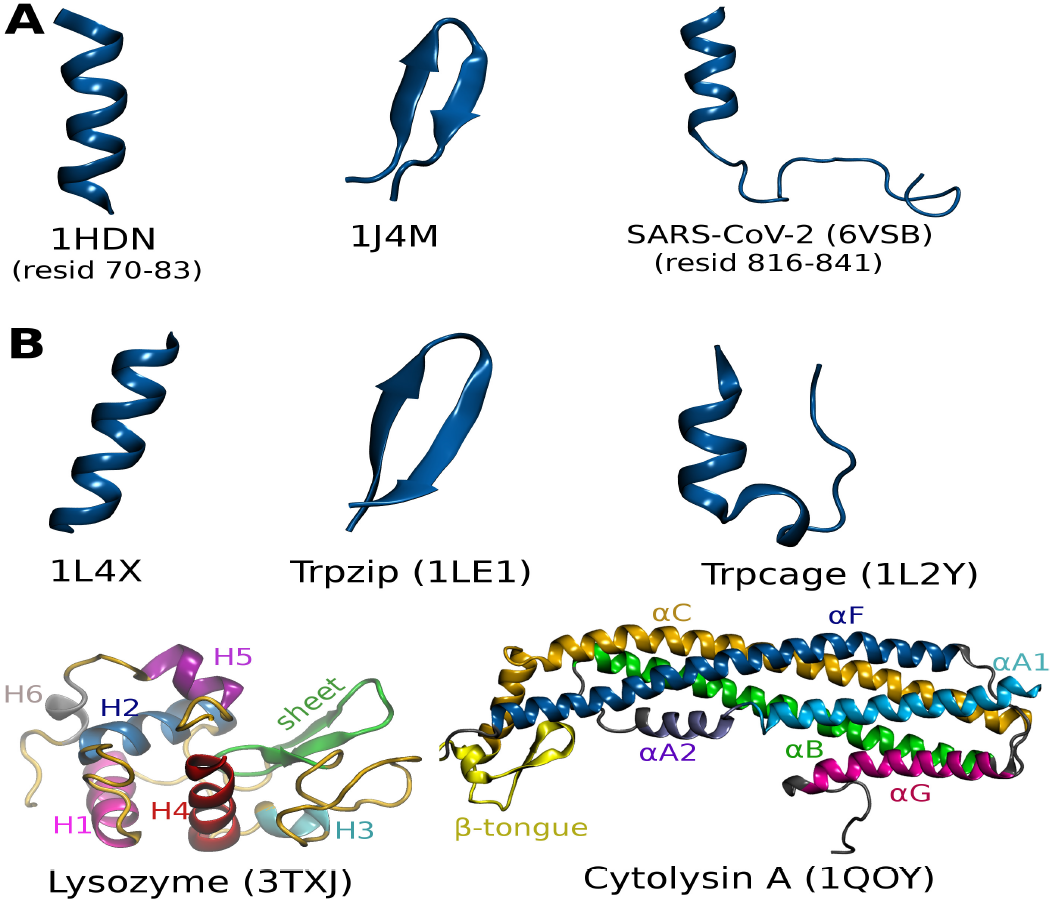
The peptides simulated for (A) parametrization and (B) validation of the CG force field are illustrated with their protein data bank (PDB) entries. Lysozyme is a multidomain protein with 129 residues comprising of helical segments, H1 (pink), H2 (blue), H3 (cyan), H4 (red), H5 (purple), H6 (silver) and extended sheet structure (green). Cytolysin A (ClyA) is a pore-forming toxin containing structural segments, namely *α*A1 (cyan), *α*A2 (violet), *α*B (green), *α*C (orange), *β*-tongue (yellow), *α*F (blue), and *α*G (pink).

### 2.1 Atomistic simulations

The peptides shown in Fig. 1 were simulated in water with the all-atom (AA) CHARMM36 force fields using the GROMACS (2018.2) MD engine. ^69,70^ The CHARMM

TIP3P water model was used for the aqueous environment, and periodic boundary conditions were used in the three orthogonal directions. The details for the different systems investigated are given in Table 1, and data was acquired over 200 ns long simulations after equilibration for 10 ns. The simulations were performed at room temperature (303.15 K) using a Nosé-Hoover thermostat with a coupling time constant of 1 ps. ^71^ While the system pressure was maintained at 1 bar using a Parrinello-Rahman barostat with a isotropic pressure coupling time constant of 5 ps and compressibility 4.5 × 10^−5^ bar^−1^, which corresponds to water compressibility at ambient conditions. ^72^ The bonds with H-atoms were constrained using the LINCS algorithm. ^73^ The equations of motion were integrated using the leap-frog algorithm with a step size of 2 fs. The non-bonded van der Waals interactions were computed within a cutoff distance of 1.2 nm, and the forces were smoothly truncated to zero over the distance from 1.0 to 1.2 nm. The long range electrostatic interactions for charged species were evaluated using the Particle Mesh Ewald (PME) method with a real-space cutoff distance of 1.2 nm. ^74^ Na^+^ and Cl^−^ ions were added to neutralize charges on the peptides.

**Table 1.**
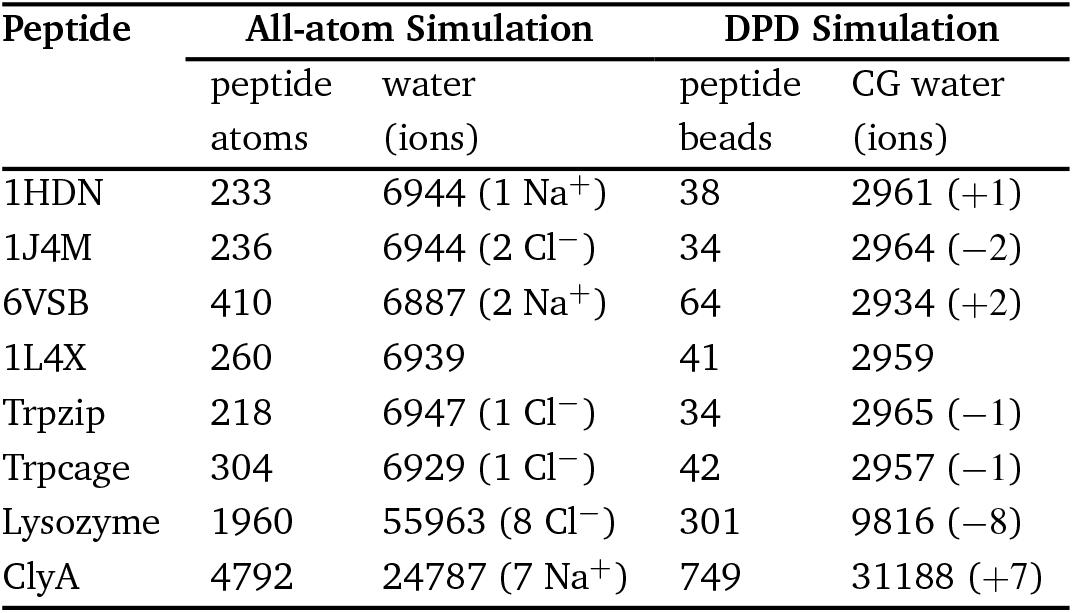
Details for different systems simulated using all-atom and DPD descriptions

### 2.2 Dissipative particle dynamics simulations

We use the standard formulation of the DPD method, ^41,42,59,75^ where suitable molecular fragments or beads represent the level of coarse-graining. The DPD beads interact with each other via the centrally directed pairwise forces. The soft repulsive conservative force *f* ^*C*^ between DPD particles *i* and *j* positioned at **r**_**i**_ and **r**_**j**_ is,

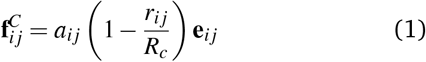

where *a*_*i j*_ is the repulsive force constant and *R*_*c*_ is the cutoff radius. The radial separation between the particles *i* and *j* is *r*_*i j*_ = |**r**_*i*_ − **r**_*j*_|, and the unit vector **e**_*i j*_ = (**r**_*i*_ − **r** _*j*_)*/r*_*i j*_. The DPD particles also interact through pairwise dissipative *f* ^*D*^ and stochastic *f* ^*R*^ forces given by,

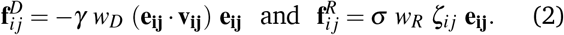

The relative velocity **v**_*i j*_ is **v**_*i*_ − **v**_*j*_, while *ζ*_*i j*_ is a random variable drawn from a Gaussian distribution with zero mean and a unit variance. While *γ* and *σ* are the strengths of the dissipative and random forces respectively. The weight function *w*_*R*_(*r*) = 1 − (*r*_*i j*_*/R*_*c*_). The strengths of the dissipative and stochastic forces are governed by the fluctuation-dissipation theorem, ^76^

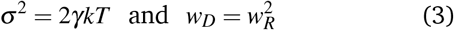

resulting in a thermostat for the DPD simulation. Accordingly, the noise amplitude *σ* = 3.0 and the frictional strength *γ* = 4.5 are the usual choice for the DPD thermostat. The energy scale is *kT*, where *k* and *T* denote the Boltzmann constant and absolute temperature, respectively. The equations of motion for the DPD particles are integrated using the velocity Verlet scheme. ^42,77^

### 2.3 Electrostatics in DPD simulations

The charges assigned to the side chains for some of the amino acids are illustrated in Fig. 2. In addition, peptides were assigned a charge of +1e at the N-terminus and -1e at the C-terminus. ^55,65^ Electroneutrality is maintained by addition of ions (see Table 1). Electrostatic interactions among the charged DPD particles can be evaluated either by locally solving the electrostatic field on spatially distributed grids ^75^ or by using the Ewald summation. ^78–80^ A comparative assessment of these methods is detailed elsewhere. ^81^ We computed the electrostatic potential ϕ by solving iteratively the Poisson equation on real-space grids, ^75^

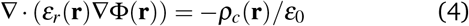

**Fig. 2.**
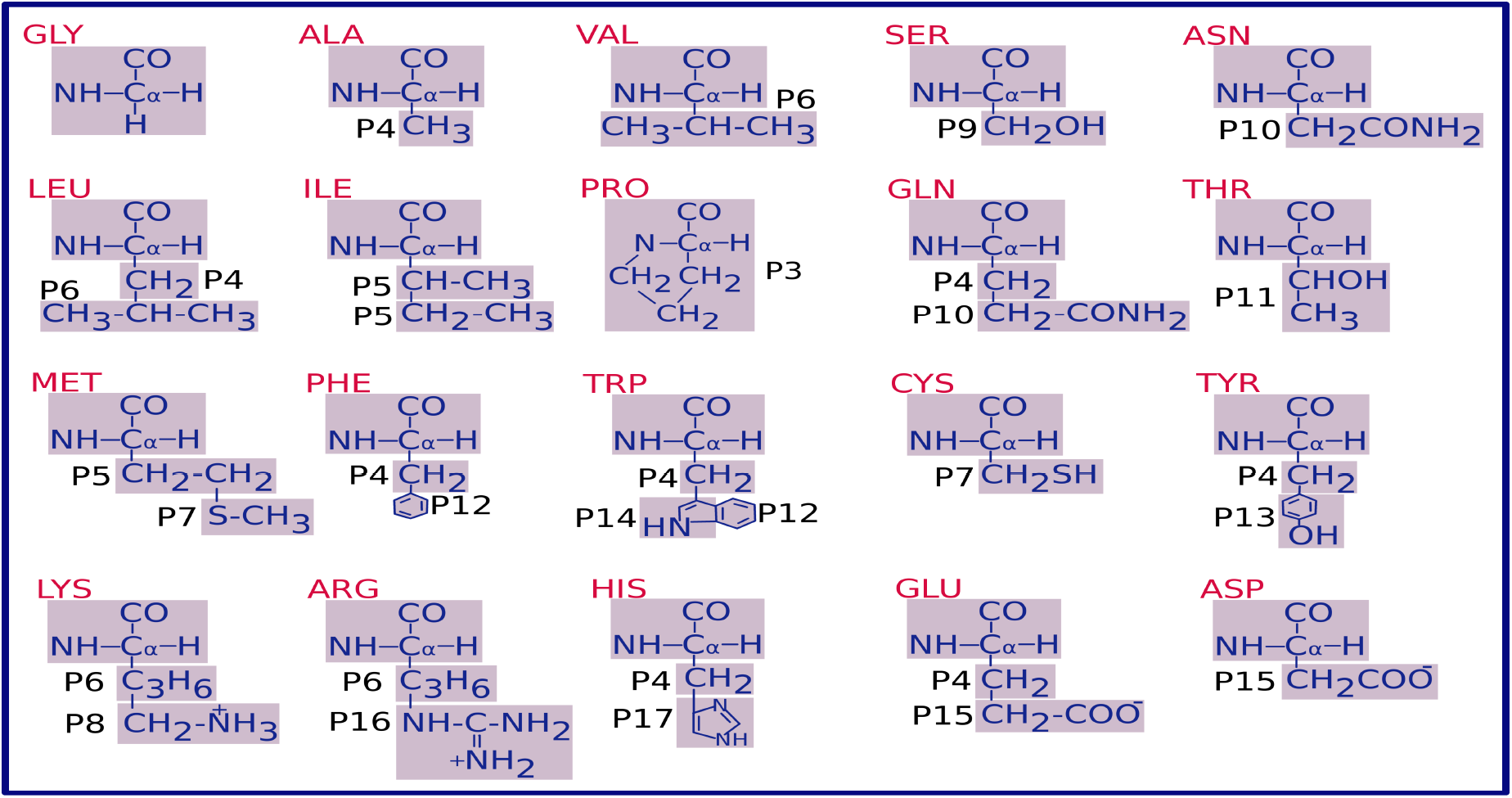
Schematic showing the mapping of amino acids onto the CG beads illustrated as shaded boxes. The backbone CG beads are centered on the C_*α*_ atoms of the amino acid residues and charges are illustrated for some of the side chains. The residues GLY and PRO are represented by a single bead. The bead type ascribed to the backbone bead is P2, except for the PRO residue, which has particle type P3. The other side chain beads have particle types, P4 - P17 as labelled in the schematic. The repulsive force parameters for the different particle types are given in Table 2.

The constants *ε*_0_ and *ε*_*r*_ are the dielectric permittivity of vacuum and local relative dielectric constant, respectively. We used a uniform dielectric constant, *ε*_*r*_ = 78. The computational domain is discretized into three dimensional cells with a mesh size *R*_*c*_. The Coulomb charges on the DPD particles are smeared out over the grid points within a distance of *R*_*e*_ = 1.6*R*_*c*_ using a linear charge distribution function,

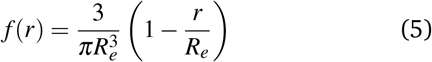

Following Groot, ^75^ the grid-based electrostatic potential, Φ(**r**) is obtained by solving the discretized version of the Poisson equation (eqn (4)) using the successive relaxation method. The electrostatic force on each particle is calculated by interpolating the gradient of the potential using the same distribution function *f* (*r*).

The system details including the number of DPD water beads are provided in Table 1. All DPD simulations are carried out at room temperature and constant volume with periodic boundary conditions in three Cartesian directions. A time step of 0.01*τ* is used for integrating the equations of motion. The time scale of *τ*∼88 ps was deduced by mapping to the diffusion coefficient of water. ^59^ In reduced DPD units the length *R*_*c*_, time *τ*, energy *kT* and masses *m* of the CG particles are set to unity. Each DPD trajectory for peptides 1HDN, 1J4M and 6VSB, which are used for parametrization, consists of 2 × 10^5^ time steps with a step size of 0.01*τ*, while each DPD simulation for 1L4X, Trpzip and Trpcage is comprised of 10^6^ time steps. The last half of the trajectories are used for analysis in each case. The DPD force field developed using the different peptides (Fig. 1) is given next.

## 3 DPD models for water and proteins

The DPD bead for representation of water effectively comprises of three water molecules with the length scale *R*_*c*_ = 0.646 nm, which is equivalent to the volume of three water molecules at room temperature. ^59^ With this three-to-one CG mapping scheme, the force constant for soft repulsive interactions between water-water is *a*_*ww*_ = 78 *kT/R*_*c*_, which corresponds to the isothermal compressibility of water at room temperature. ^59^

Amino acids are the building blocks of peptides and proteins, whose secondary structures are highly correlated with the backbone dihedral angles. ^82–84^ Therefore, in the reduced representation shown in Fig. 2, we map the backbone atoms of the protein onto the CG backbone beads (B), with the centers of these beads located at the positions of the C_*α*_ atoms. ^65^ The mapping scheme we follow in fragmenting the side chains into CG beads allows us to directly adopt the Flory-Huggins (*χ*) parameters used by Truszkowski *et al*. ^56^ for the interactions between the CG beads. The different bead types for the amino acids are labelled from P2 - P17 as illustrated in Fig. 2 where the backbone bead ‘B’ corresponds to the bead type P3 for PRO and P2 for all other residues. For notational convenience we refer to the side chain beads as ‘S1’, ‘S2’ and ‘S3’ labelled progressively outward from the backbone bead. The residues GLY and PRO consist only of backbone beads, while TRP with bulky aromatic groups is represented by one backbone bead and three side chain beads. For example P4, P14 and P12 in the TRP residue represent the side chain beads S1, S2 and S3, respectively, while the remaining 17 amino acid residues have either one (S1) or two (S1 and S2) side chain beads. The bonded interactions between the different beads are modelled using harmonic potentials with appropriate spring constants,

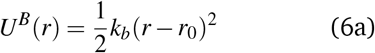

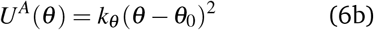

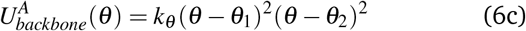

where *U*^*B*^(*r*) is the bonded potential with equilibrium bond length *r*_0_, *U*^*A*^(*θ*) is the angle bending potential with equilibrium angle *θ*_0_, while the corresponding spring constants are *k*_*b*_ and *k*_*θ*_ respectively. Here 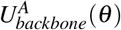 represents a dual-basin angle potential (eqn (6c)) between three consecutive beads (B-B-B) along the backbone with the equilibrium angles *θ*_1_ = 90° and *θ*_2_ = 120°, which correspond to the backbone angles for *α*-helix and *β*-sheet structures, respectively. This parametrization is an improvement and a departure from the existing DPD force fields reported in the literature. Additionally we incorporate the backbone dihedral potential,

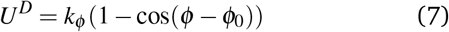

where the torsion angles along the protein backbone are maintained at the native state value of *ϕ*_0_ using a cosine potential with the stiffness constant *k*_*ϕ*_. The dihedral potential has recently been used to stabilize the *α* helical structure in polyalanine. ^58^

## 4 Model parametrization

The repulsive force constants *a*_*i j*_ among the protein beads and protein-water interactions are computed using the Flory-Huggins energy parameters *χ*_*i j*_ using, ^42^

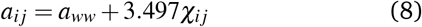

As mentioned earlier, we have adopted the Flory-Huggins parameters for the DPD beads of proteins from the work of Truszkowski *et al*., ^56^ where molecular fragments were simulated in an aqueous solvent using the COMPASS force field. ^85^ The full interaction matrix of the repulsive force constant evaluated from eqn (8) is tabulated in Table 2, where the *P*_*i*_’s represent the different bead types as described in Fig. 2. We note that ions interact with the protein and other CG beads with the same repulsive strength as the CG water, and hence ions are assigned P1 type bead. Since many DPD models for lipids membranes use a repulsive force with *a*_*ww*_ = 78 *kT/R*_*c*_, our model for the protein can directly be integrated with the existing phospholipid force fields to investigate protein-membrane complexes. ^53,54,59,86–89^

**Table 2.** Repulsive force constants, *a*_*i j*_ evaluated from eqn (8) for interactions between the different protein beads, P2-P17 as illustrated in Fig. 2 as well the water bead, P1

| Bead type → | P1 | P2 | P3 | P4 | P5 | P6 | P7 | P8 | P9 | P10 | P11 | P12 | P13 | P14 | P15 | P16 | P17 |
| --- | --- | --- | --- | --- | --- | --- | --- | --- | --- | --- | --- | --- | --- | --- | --- | --- | --- |
| ↓ |  |  |  |  |  |  |  |  |  |  |  |  |  |  |  |  |  |
| P1 | 78 | 78 | 78 | 91 | 93 | 94 | 90 | 73 | 70 | 77 | 74 | 91 | 76 | 83 | 68 | 77 | 76 |
| P2 |  | 78 | 72 | 80 | 77 | 77 | 76 | 72 | 75 | 71 | 74 | 75 | 71 | 72 | 72 | 75 | 72 |
| P3 |  |  | 78 | 78 | 75 | 74 | 73 | 73 | 73 | 72 | 72 | 73 | 67 | 69 | 69 | 74 | 71 |
| P4 |  |  |  | 78 | 73 | 74 | 73 | 80 | 82 | 81 | 83 | 76 | 80 | 78 | 96 | 101 | 81 |
| P5 |  |  |  |  | 78 | 72 | 72 | 78 | 82 | 78 | 81 | 73 | 78 | 75 | 95 | 100 | 79 |
| P6 |  |  |  |  |  | 78 | 72 | 78 | 82 | 78 | 81 | 72 | 77 | 74 | 97 | 101 | 79 |
| P7 |  |  |  |  |  |  | 78 | 76 | 81 | 77 | 80 | 71 | 75 | 73 | 92 | 93 | 77 |
| P8 |  |  |  |  |  |  |  | 78 | 72 | 71 | 73 | 77 | 69 | 72 | 63 | 68 | 73 |
| P9 |  |  |  |  |  |  |  |  | 78 | 75 | 72 | 81 | 72 | 78 | 66 | 71 | 73 |
| P10 |  |  |  |  |  |  |  |  |  | 78 | 75 | 76 | 72 | 73 | 72 | 73 | 73 |
| P11 |  |  |  |  |  |  |  |  |  |  | 78 | 80 | 71 | 76 | 67 | 73 | 73 |
| P12 |  |  |  |  |  |  |  |  |  |  |  | 78 | 75 | 72 | 93 | 94 | 76 |
| P13 |  |  |  |  |  |  |  |  |  |  |  |  | 78 | 73 | 71 | 72 | 68 |
| P14 |  |  |  |  |  |  |  |  |  |  |  |  |  | 78 | 82 | 80 | 71 |
| P15 |  |  |  |  |  |  |  |  |  |  |  |  |  |  | 105 | 56 | 70 |
| P16 |  |  |  |  |  |  |  |  |  |  |  |  |  |  |  | 105 | 74 |
| P17 |  |  |  |  |  |  |  |  |  |  |  |  |  |  |  |  | 78 |

Using a bottom-up approach for coarse-graining, the bonded parameters, namely the equilibrium bond lengths and angles (*r*_0_ and *θ*_0_ in eqn (6)), are deduced from the centers of reference distributions for the virtual CG bonds and angles derived from AA trajectories. We used an iterative procedure to optimize the bonded interactions. In order to use a higher time step in DPD simulations, the bond stiffness *k*_*b*_ is restricted to 2000 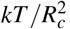, and a minimum value of *k*_*b*_ is set to 1500 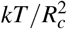 to control the spread of the distributions. The stiffness for the angle bending potential is chosen to be 10, 15 or 20 *kT/rad*^2^ depending on the spread of the angles in the reference distributions. During the iterative procedure, bond lengths (*X* = *r*_0_) as well as the angles (*X* = *θ*_0_) in CG distributions (*X*^*CG*^) are updated, according to *X* = *X* + (*X*^*AA*^ - *X*^*CG*^), with a convergence tolerance of 0.005 nm in bond lengths and a maximum 5° deviation in angles from the centers of corresponding reference distributions (*X*^*AA*^).

The optimized CG bond lengths and the bond stiffness values are given in Table 3. With *R*_*c*_ = 0.646 nm, the CG bond length of 0.58*R*_*c*_ between two consecutive backbone beads (B-B) results in a distance of 0.38 nm between two consecutive C_*α*_ atoms in the AA protein model. The bond between S1 and S3 beads of TRP is also constrained to 0.64*R*_*c*_ for controlling angles in the TRP residue.

**Table 3.** Bonded interaction parameters between the backbone beads, B and the side chain beads S1, S2 and S3 for the different amino acid residues illustrated in Fig. 2

| Residue | Bonds<br>(B-S1, S1-S2) |  | Angles<br>(B-S1-S2) |  |
| --- | --- | --- | --- | --- |
| | bond length $r_0$<br>($R_c$) | stiffness $k_b$<br>($\text{kT}/R_c^2$) | angle $\theta_0$<br>(degree) | stiffness $k_\theta$<br>($\text{kT}/\text{rad}^2$) |
| ALA | 0.24 | 2000 | - | - |
| ARG | 0.38, 0.56 | 1500, 1500 | 140 | 10 |
| ASN | 0.40 | 1500 | - | - |
| ASP | 0.39 | 1500 | - | - |
| CYS | 0.37 | 1500 | - | - |
| GLN | 0.24, 0.41 | 2000, 1500 | 90 | 15 |
| GLU | 0.24, 0.40 | 2000, 1500 | 140 | 15 |
| HIS | 0.24, 0.43 | 2000, 1500 | 110 | 20 |
| ILE | 0.31, 0.39 | 1500, 1500 | 80 | 20 |
| LEU | 0.24, 0.33 | 2000, 1500 | 120 | 20 |
| LYS | 0.37, 0.48 | 1500, 1500 | 140 | 15 |
| MET | 0.31, 0.40 | 1500, 1500 | 110 | 15 |
| PHE | 0.24, 0.48 | 2000, 1500 | 110 | 20 |
| SER | 0.31 | 1500 | - | - |
| THR | 0.31 | 1500 | - | - |
| TRP | 0.24, 0.43 | 2000, 1500 | 115 | 20 |
| TRP | 0.35 (S2-S3) | 1500 | 108 | 20 (S1-S2-S3) |
| TRP | 0.64 (S1-S3) | 1500 | - | - |
| TYR | 0.24, 0.55 | 2000, 1500 | 110 | 20 |
| VAL | 0.31 | 1500 | - | - |
| Backbone | 0.58 (B-B) | 1500 | 115 | 15 (B-B-S1) |

Figure 3 shows the distributions for bonds in the different amino acid residues compared with the corresponding target distributions obtained from atomistic simulations. The CG distributions obtained in DPD simulations are inherently broader due to the soft repulsive nature of the DPD interactions, however excellent overlap is observed with the mean values coinciding well with the AA target distributions. For residues LYS and ILE, the bond distributions are bimodal in the AA simulations, while the harmonic bond potential in DPD captures one of the peaks. Figure 4 indicates the distributions of angles within the different amino acid residues, namely B-S1-S2, as well as the angle B-B-S1 across two consecutive residues. The B-B-S1 angle is set to 115°. The angle S1-B-B (not shown) comprising of first two residues at the N-terminus is constrained to *θ*_0_ = 100° with the stiffness constant *k*_*θ*_ = 15 *kT/rad*^2^. On account of having three CG beads in its side chain, TRP residue has an additional angle S1-S2-S3 peaked at ∼108°. According to eqn (6c), the energy landscape for the backbone angle B-B-B captures dual basins, 90° and 120°, encompassing the *α*-helices as well as *β*-extended structures which will be illustrated later in the text. The stiffness constant *k*_*θ*_ is set to 15 *kT/rad*^4^ for the B-B-B backbone angle. Table 3 provides the angle bending parameters.

**Fig. 3.**
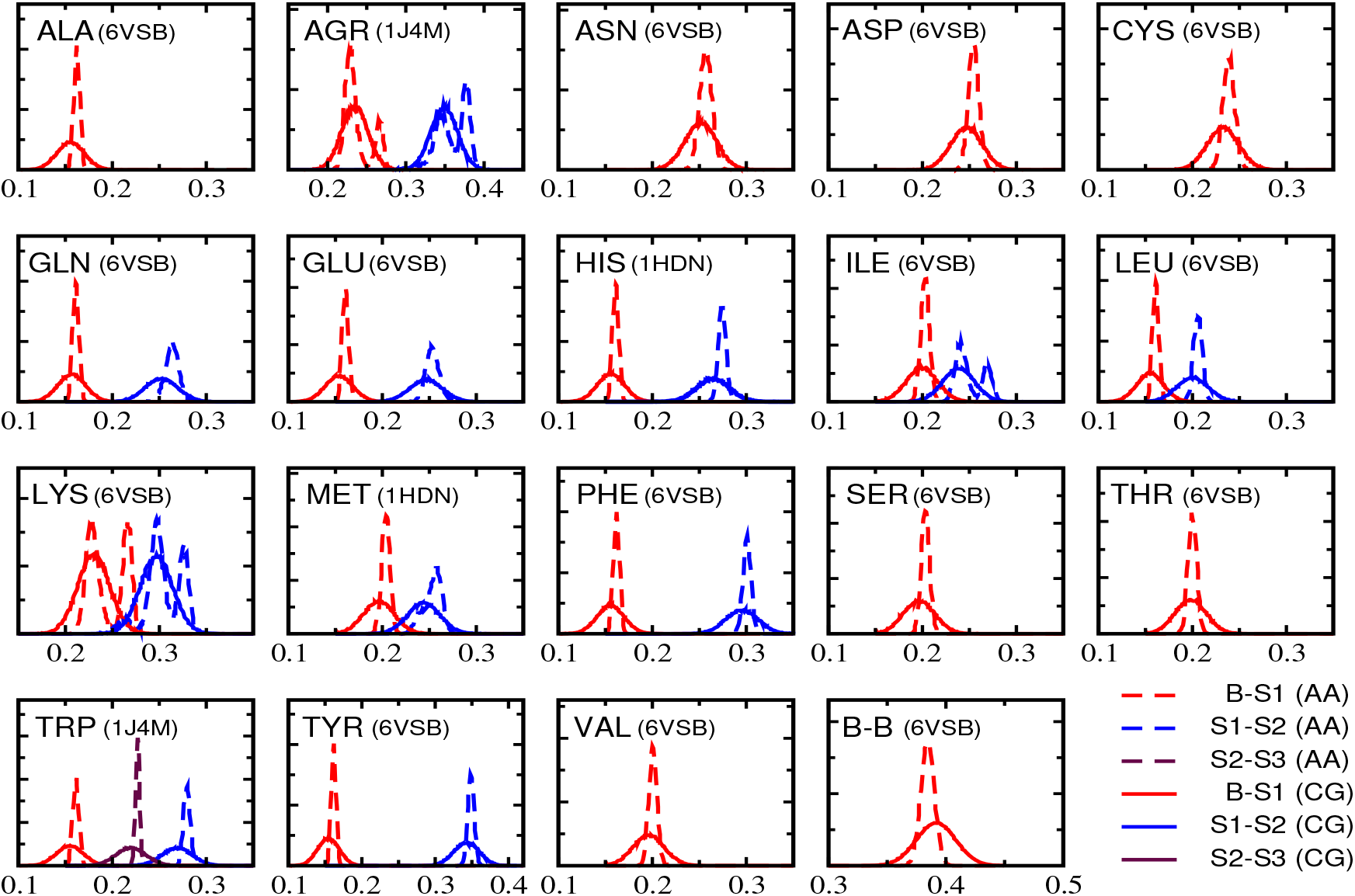
Distributions for bond lengths in amino acid residues. The solid and dotted curves indicate the distributions obtained in AA and DPD simulations, respectively, for peptides 1HDN, 1J4M and 6VSB. The bond B-S1 (red) refers to the bond between the backbone bead B and side chain bead S1, while S1-S2 (blue) is the bond between S1 and S2 beads in the side chains. The TRP residue comprises of three beads in side chain, and thus has an additional S2-S3 bond (maroon). The distribution along the peptide backbone is B-B (red). The scale on the x-axis is the bond length in nm.

**Fig. 4.**
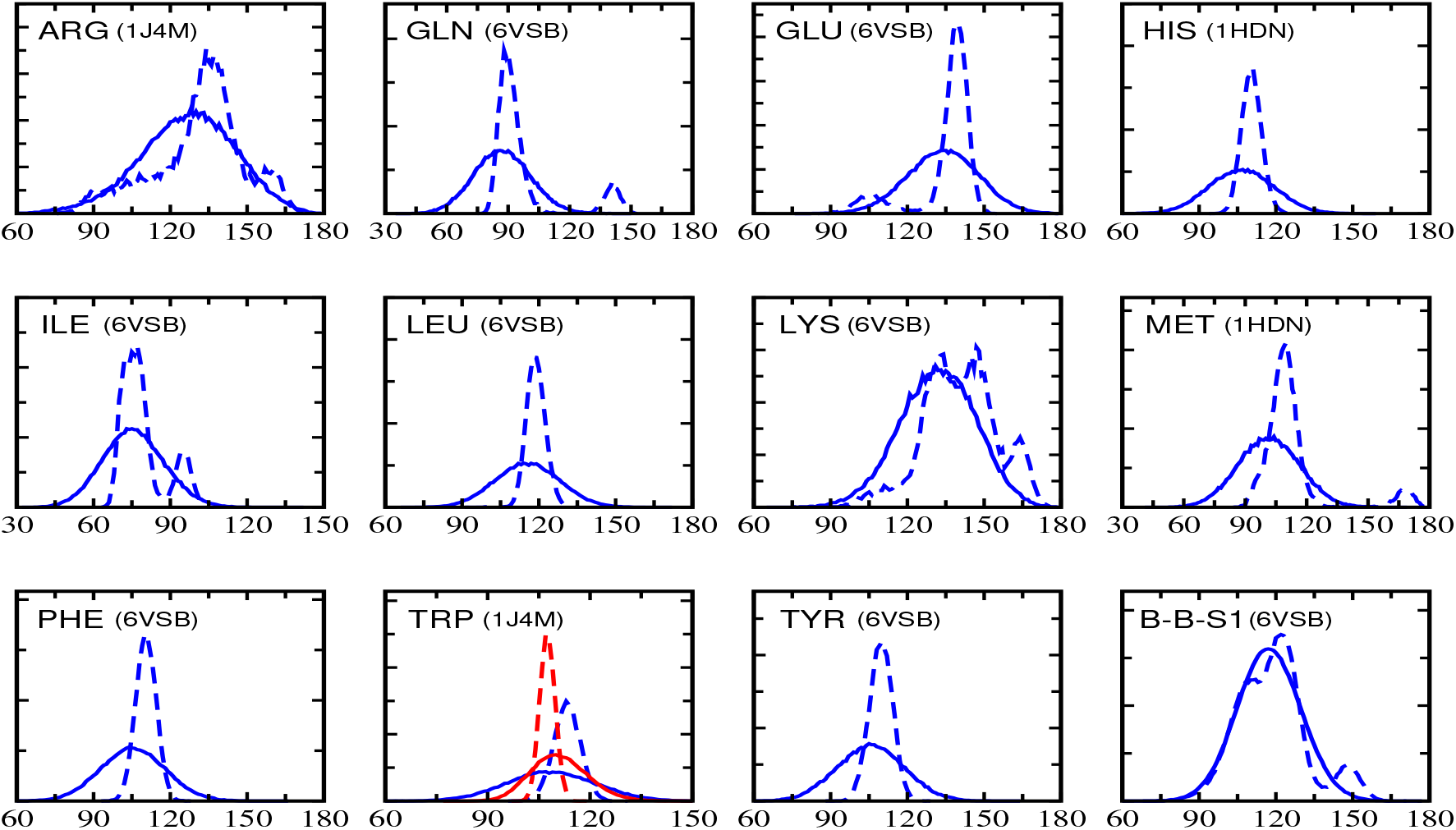
Distributions for angles in amino acid residues obtained from AA (−−) and DPD (−) simulations of peptides 1HDN, 1J4M and 6VSB. Within amino acids, there are B-S1-S2 (blue) and S1-S2-S3 (red in TRP) angles. The distribution for the angle across two consecutive residues, B-B-S1, is obtained by averaging over amino acids in a peptide.

Stabilization of the native state conformations of peptides is achieved through the backbone dihedrals. The torsion angles *ϕ*_0_ between four consecutive C_*α*_ atoms along the protein backbone are computed from the native crystal structure, and maintained in DPD simulations using the torsion potential (eqn (7)) with the stiffness constant *k*_*ϕ*_ = 10*kT*. The values for *ϕ*_0_ are provided in Table S1 in the Electronic Supplementary Information (ESI).

## 5 Model validation

The proposed DPD parameters for the protein simulations are validated by evaluating various protein structural metrics such as the root mean square deviation (RMSD), radius of gyration (*R*_*g*_), residue-residue contacts and the Ramachandran-like maps. The proposed DPD force field is validated using the peptides 1L4X (*α*-helix), Trpzip (*β*-hairpin) and Trpcage (helix+loop), where we make a direct comparison with the atomistic simulations thereby providing a stringent test to the accuracy of the proposed force field.

### 5.1 Structural similarity

To measure the similarity between the protein conformations and its native structure, we have calculated the root mean square deviation, RMSD(**r, r**′) = 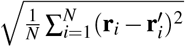, where *N* denotes the number of amino acid residues in the peptide sequence. While **r** and **r**′ are position vectors of backbone beads (C_*α*_ atoms for AA model) in the simulated trajectory and in the native PDB structure, respectively. The ability of the DPD model to retain the level of compactness of the protein structure close to that of the native structure is determined by computing the radius of gyration, 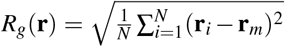, where **r**_*m*_ is the center of mass coordinates of the protein backbone.

Figure 5 shows the joint probability histograms of RMSD and *R*_*g*_ for the peptides 1L4X (*α*-helix), Trpzip (*β*-hairpin) and Trpcage (helix+loop). The color maps are represented as -log(*P/P*_*max*_) where *P*_*max*_ is the maximum value of *P* in the distribution. Hence the intensity value close to zero indicates the highest probabilities in the distribution. From the high probability regions the radius of gyration is accurately captured in our DPD simulations. For instance, the distribution for the peptide Trpzip peaks at *R*_*g*_∼0.6 ± 0.001 nm in DPD model (Fig. 5E), and this agrees well with *R*_*g*_∼0.58 ± 0.001 nm at the high probability region for the AA model (Fig. 5B).

**Fig. 5.**
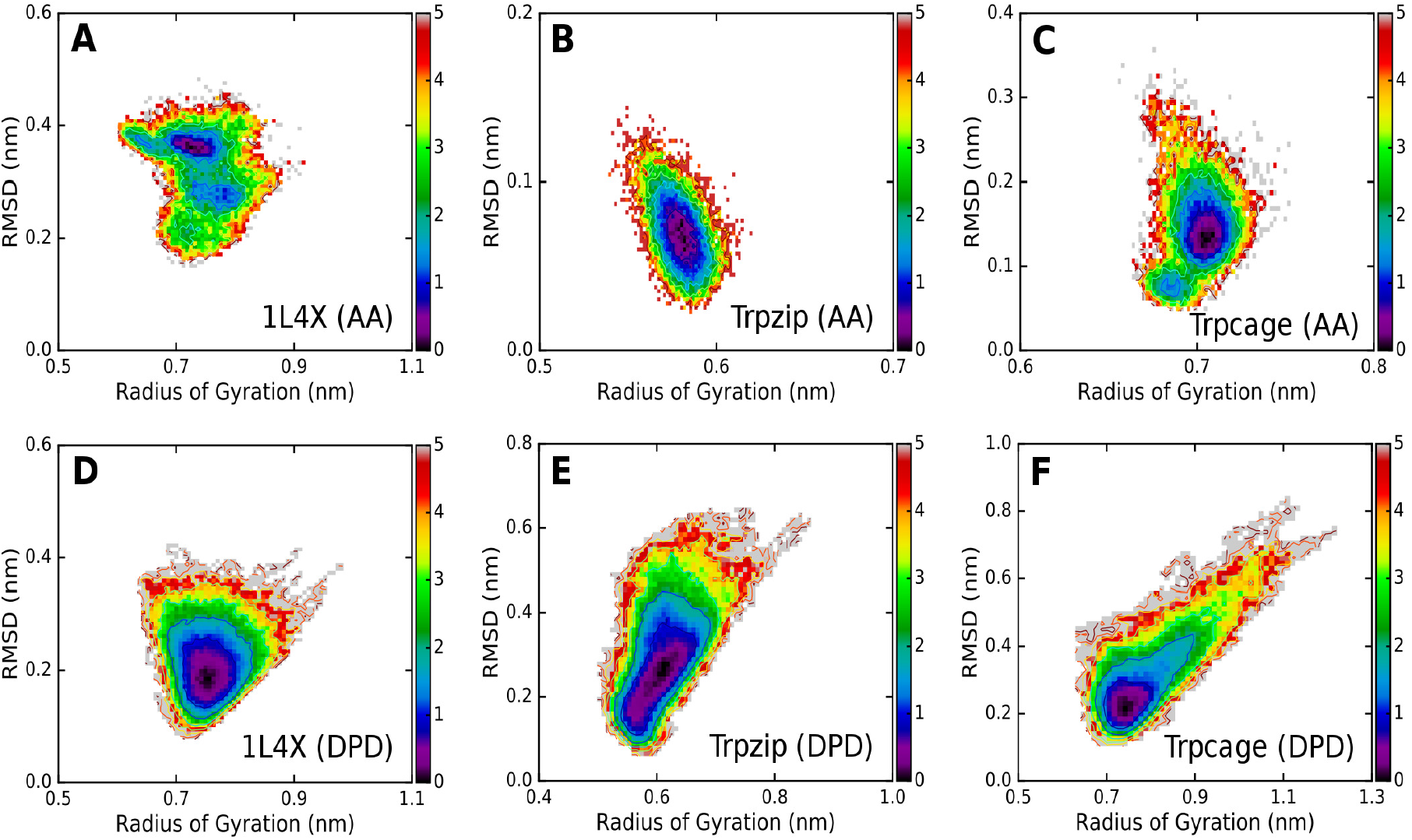
Color maps for the joint probability histograms *P* for the root mean square deviation (RMSD) and radius of gyration on the log-normal scale of -log(*P/P*_*max*_), where *P*_*max*_ is the maximum value of *P* in histograms. The histograms (A, B, C) refer to the AA simulations for peptides 1L4X (A), Trpzip (B) and Trpcage (C), while their DPD counterparts are shown in panels D, E, and F respectively.

The RMSD variations show some interesting trends across the different proteins. For the *α*-helical peptide 1L4X, the range of the RMSD variations are similar, with the DPD model displaying a slightly lower RMSD when compared with the AA data. In the case of Trpzip (*β*-hairpin) and Trpcage (helix+loop), the RMSD values obtained from the AA simulations are smaller when compared with the DPD simulations. Although the RMSD of the peptides in the DPD simulations shows a slightly higher spread when compared to a relatively narrow range observed in the AA simulations, the conformations sampled at RMSD ranging from ∼0.1 − 0.3 nm represent the energetically favourable configurations that are sampled to a greater extent (Fig. 5). Furthermore, if one examines the most probable values for RMSD, the differences are within 0.2 nm between DPD results and the reference AA RMSD data. Additionally, the range of the RMSD sampling is superior to the previous polarizable model, ^55^ which reported RMSD in the range 0.5 − 0.7 nm for the peptides Trpzip and Trpcage. For the peptides 1HDN, 1J4M and 6VSB, which were used for parametrization, the RMSD and *R*_*g*_ data are provided in Fig. S1 of the ESI.

### 5.2 Native contacts

The success of CG models for proteins relies on its ability to retain the residue-residue contacts that characterize a given native structure. This can be achieved by constraining the contacts using the harmonic potential (Gō and elastic network models) or through Lennard-Jones potential (Gō-MARTINI). ^17,38,57,90^ In the DPD model presented here, the secondary structures are stabilized with the dihedral (eqn (7)) and the dual-basin angle bending potentials (eqn (6c)). Unless stated otherwise, we computed the contact maps using a distance criterion between a pair of residues falling within 1 nm for both the AA and DPD simulations.

As illustrated in Fig. 6, where contact maps with the color bars indicate the fraction of a trajectory having stabilized contacts, the residue-residue contacts are well captured in the helical peptides (Fig. 6A and D). The comparison for the peptides with *β*-hairpin (Fig. 6B and E) and the flexible loop structures (Fig. 6C and F) are also satisfactory. The discrepancies in contact maps between the AA and DPD models for these structures are attributed to non-local excluded volume interactions, which are soft repulsive in the DPD force field (eqn (1)). The soft repulsive nature of the DPD force field is able to capture all the dominant contact pairs, albeit with reduced intensity in some cases. For example in the Trpzip structure (Fig. 6B and E)) the contacts between residues 12 with 1-4 are captured with a lower intensity in the DPD simulations. Although an increased cutoff for the contacts in the DPD model can reduce the differences, we prefer to make comparisons with the AA simulations with a similar cutoff criterion. Figure S2 in ESI provides the contact maps for the peptides 1HDN, 1J4M and 6VSB. The native contacts are further quantified by computing the order parameter ^91^ 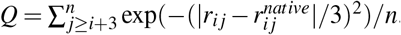. The summation runs over all non-local contacts between pairs of residues *i* and *j* ≥ *i* + 3, with distances *r*_*ij*_ in simulated trajectories and 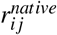 in native PDB structures. The order parameter *Q* is normalized by the total contacts, *n*. The time averaged *Q* values from the DPD simulations are 0.79, 0.63 and 0.66 for 1L4X, Trpzip and Trpcage, respectively (Fig. S3 in ESI). The Q values lying in the range 0.55 ≤ *Q* ≤ 1 provides additional confirmation for the inherent stability of the structures in the proposed DPD force field.

**Fig. 6.**
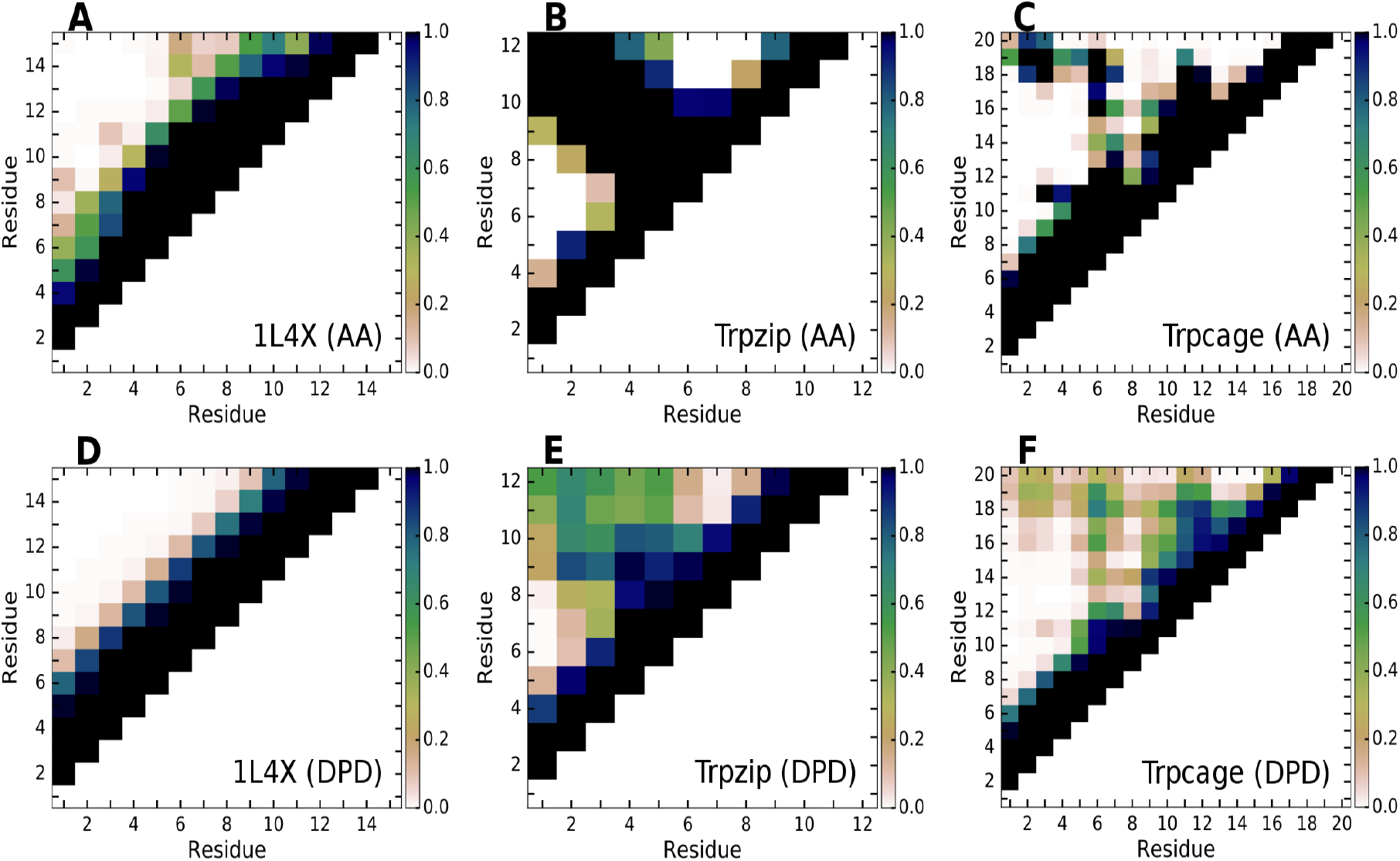
The residue-residue contacts with a cutoff distance of 1.0 nm are shown for the peptides 1L4X (A, D), Trpzip (B, E) and Trpcage (C, F). The contact maps obtained from the all-atom simulations are indicated in panels A, B, and C, while those derived from the DPD simulations are shown in panels D, E and F.

### 5.3 Ramachandran conformational space

Equivalent to an energy surface on the *ϕ* − *ψ* plane formed by two dihedral angles in atomistic models, the Ramachandran-like plot in a reduced CG representation is a two-dimensional density plot of the backbone angle *θ* and torsion angle *ϕ*. ^92^ The Ramachandran-like maps for peptides 1HDN, 1J4M and 6VSB are given in Fig. S4 of ESI. Referring to Fig. 7A and D, dominant sampling of the conformations occurs at, *θ* = 90° and *ϕ* = 50° for the helical peptide 1L4X as well as for the peptide Trpcage (Fig. 7C and F), which has helices at the N-terminus (residues 1-9). The increased sampling centered around *ϕ* = −120^*°*^ accounts for the turn regions in the peptides Trpzip (Fig. 7B and E) and Trpcage (Fig. 7C and F). The presence of *β*-sheets in the peptide Trpzip (Fig. 7B and E) is evident from high density spots at *ϕ* ∼±180°. Correlating the conformational space mapped with the *ϕ, θ* angles with the observed secondary structures, our data is consistent with the density maps in these reduced co-ordinates given in earlier CG-MD studies. ^92^

**Fig. 7.**
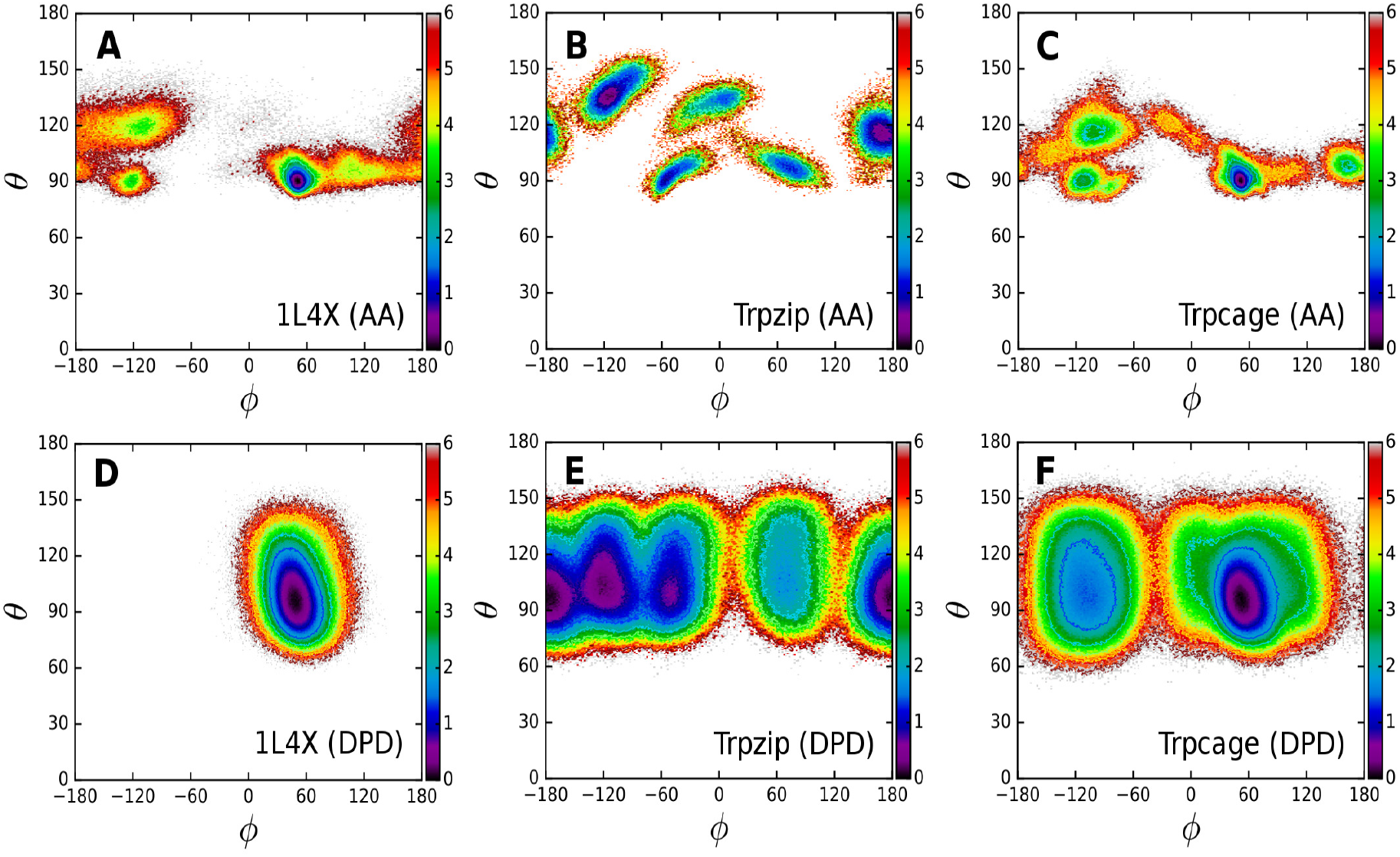
Ramachandran-like plots for the model peptides 1L4X (A, D), Trpzip (B, E) and Trpcage (C, F), showing in color maps the probability density *P*(*ϕ, θ*) as -log(*P*(*ϕ, θ*)*/P*_*max*_) on phase space of backbone angle *θ* and dihedral angle *ϕ*. The maximum density is *P*_*max*_. The top panels (A, B, C) are constructed from AA simulations, while the panels at the bottom (D, E, F) are the results from the DPD simulations. The angles are represented in degrees.

### 5.4 Protein folding

The robustness of the proposed force field is tested by allowing the peptide sequences to fold back to their three dimensional native structures. We carried out folding simulations on peptides 1L4X (*α*-helix) and Trpzip (*β*-hairpin). The initial unfolded configurations were obtained by switching off the dihedral terms and changing the equilibrium backbone angles to 180°. With ‘d’ being the distance between *C*_*α*_ atoms in the native state in the AA model, a pair of residues in the DPD model is said to be in contact when their backbone beads are closer than 1.3d. ^90^ However, due to purely repulsive nature of the DPD forces and relatively higher flexibility of *β*-hairpin in absence of hydrogen bonding, a cutoff of 1.3d + 0.2 nm can be used to capture more than 90% of the native contacts in *β*-hairpin structure. Figure 8 illustrates the time evolution of native contacts averaged over 10 independent folding simulation trajectories for each peptide. It is clear from the folding curves in Fig. 8 that the characteristic folding times for *α*-helix as well as *β*-hairpin are of similar order (∼ 10*τ*), which we attribute to the equilibration time scales for the dihedral angles. Unlike the ten-fold faster folding of *α*-helix over the *β*-hairpin structures observed for atomistic as well as Gō-MARTINI models in previous studies, ^90,93^ the folding times for the *α*-helix is only slightly faster than that of *β*-hairpin in the present DPD model.

**Fig. 8.**
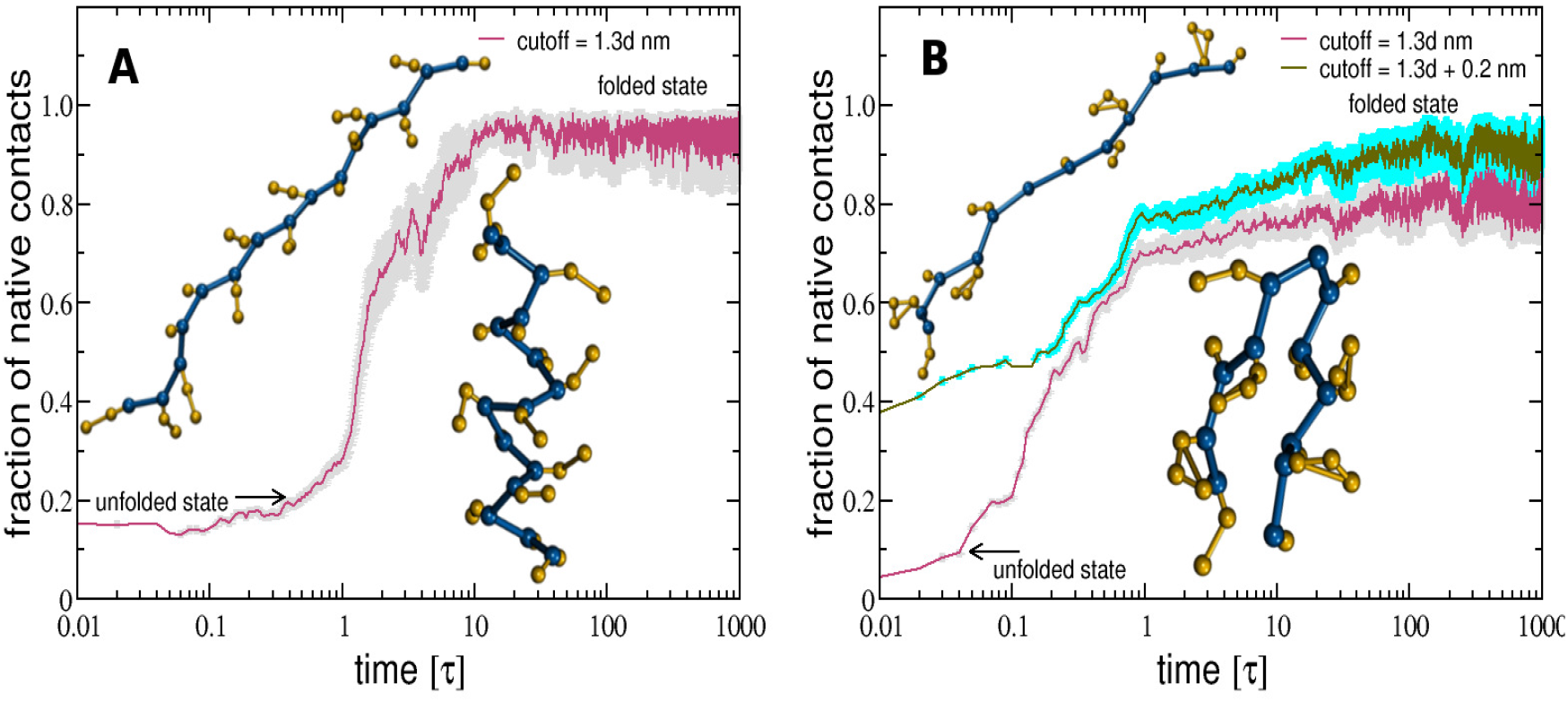
Folding of peptides (A) *α*-helix 1L4X, and (B) *β*-hairpin Trpzip to their native states. The insets show the initial unfolded and the final folded structures. The backbone beads are colored in blue, while the side chain beads are illustrated in orange. The contacts between the backbone beads were calculated using a distance cutoff of 1.3 times the distance ‘d’ between C_*α*_ atoms in the native AA structure. A slightly higher cutoff of 1.3d + 0.2 nm allows *β*-hairpin to recover more than 90% of the native contacts.

### 5.5 Multidomain proteins

The generality of the proposed DPD protein model for in-silico study of multidomain proteins is further examined by simulating larger protein structures. We chose lysozyme and cytolysin A (ClyA) for this purpose, and describe simulations first for lysozyme followed by ClyA in this section. The simulations are carried out for 10^6^ time steps after an initial equilibration period of 10^5^ steps, and the last half of the trajectory was used for calculating the structural properties. Lysozyme (PDB: 3TXJ) is employed here to illustrate the manner in which we incorporate the disulfide linkages present between pairs of CYS residues. Lysozyme comprises of 129 amino acid residues, with six small helical domains, H1 (residues:5-15), H2 (25-36), H3 (80-83), H4 (88-102), H5 (109-115), H6 (120-124), as well as extended structures (residues: 43-45, 51-53, 58-59) as depicted in Figs. 1 and 9D. The four pairs of CYS residues, (residues 6 and 127, 30 and 115, 64 and 80, 76 and 94) are involved in forming four disulfide linkages in lysozyme. In order to incorporate the disulfide bonds in the DPD model, we used a harmonic potential (eqn (6a)) between these pairs of residues with a bond length of *r*_0_ = 0.31*R*_*c*_ (0.2 nm in AA reference model) and a bond stiffness, *k*_*b*_ = 1500 *kT/nm*^2^.

**Fig. 9.**
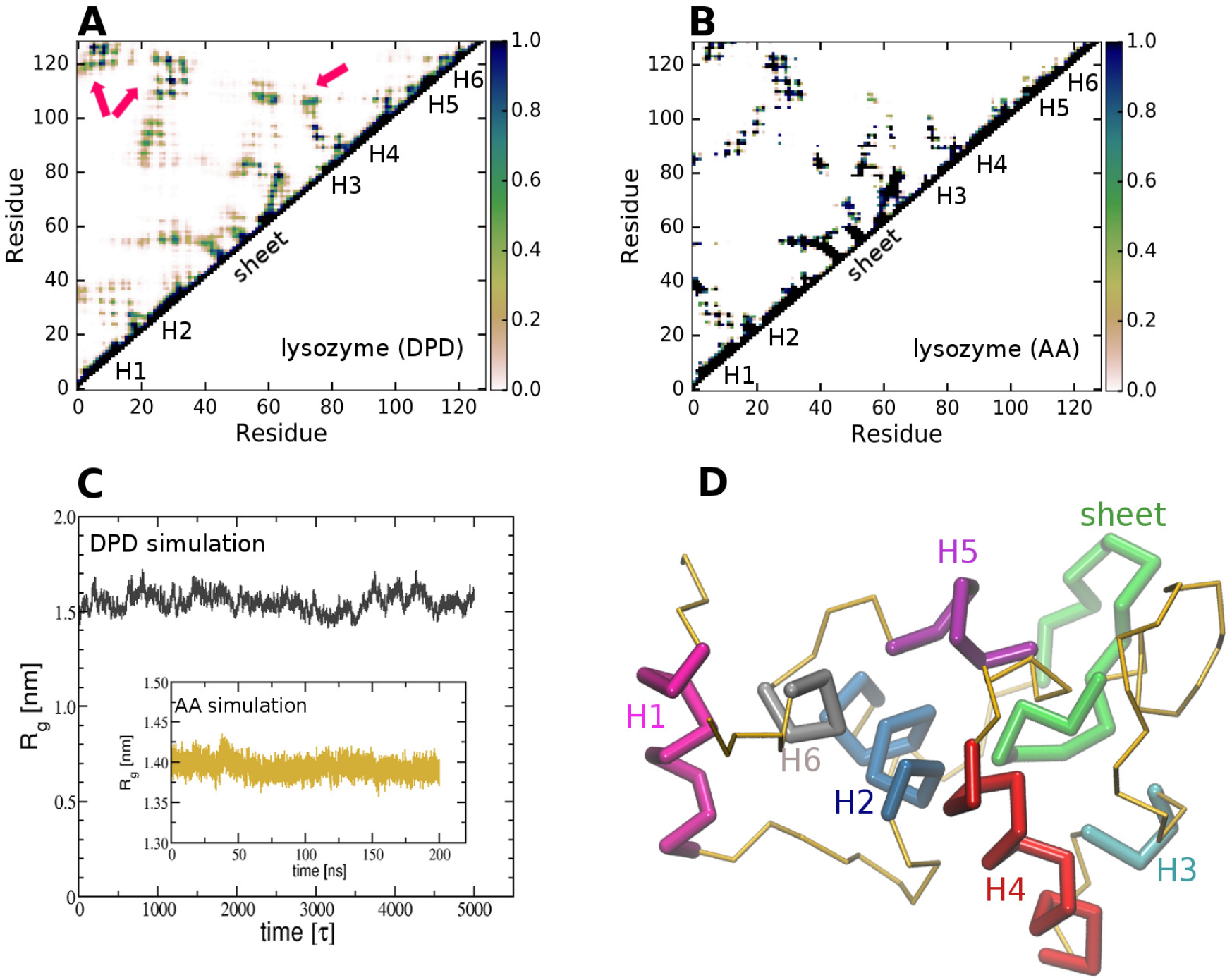
Lysozyme simulation results. (A) The residue-residue contact map in DPD simulation with cutoff distance of 1 nm, showing the contacts especially in the structural regions, namely six helices H1-H6 and the extended sheet. The arrows indicate the non-local contacts in the vicinity of the disulfide linkages. (B) The contact map obtained from AA simulation of lysozyme with a distance cutoff of 1 nm. (C) The radius of gyration (*R*_*g*_) in DPD simulation stabilizes at *R*_*g*_∼1.5 nm, while it is quite stable at *R*_*g*_∼1.4 nm in AA simulation shown in the inset. The time in DPD simulations is in units of *τ*. (D) The simulation snapshot of lysozyme is illustrated with structural segments, and the color scheme is similar that used for lysozyme in Fig. 1.

The time evolution of RMSD of the structural domains of lysozyme (Fig. S5 in ESI) indicates that the six helices and the extended strands retain their native structures at RMSD ∼0.3 nm in the DPD simulation. Due to the presence of flexible loops and turns, the overall structure of entire lysozyme showed a higher RMSD of 0.937±0.003 nm when compared with RMSD 0.117 ± 0.003 nm obtained in the atomistic simulation. However, the RMSD observed in our simulation is nearly 30% lower than the RMSD of lysozyme reported earlier. ^58^ Interestingly, as indicated in Fig. S5 in ESI, the root mean square fluctuations (RMSF data) in DPD simulation largely follow the trends observed in RMSF data obtained from AA simulation of lysozyme, lying well within ∼0.5 nm for most of the regions within the protein.

Figure 9A shows the residue-residue contacts within a cutoff distance of 1 nm in lysozyme. Our DPD force field efficiently stabilizes the residue contacts in all six helical as well as extended sheet regions of the lysozyme when compared with the contact maps derived from atomistic simulation (Fig. 9B). Furthermore, the contacts between residues forming the disulfide linkages are captured both in AA as well as in DPD results. For instance, the contacts around the pairs of residues, namely 6 and 127, 30 and 115, 64 and 80, 76 and 94, are captured in our DPD simulations. The *R*_*g*_ value of 1.549 ± 0.001 nm from DPD simulation is in good agreement with the *R*_*g*_ = 1.393 ±0.001 nm obtained from the AA simulation (Fig. 9C).

ClyA is an *α* pore-forming toxin secreted by *Escherichia coli*, which assembles on the target membrane to form a dodecameric transmembrane channel. ^94,95^ The structure of the ClyA monomer (PDB: 1QOY, 303 residues in Fig.1) is predominantly *α* helical containing the segments *α*A1 (residues 1-35), *α*A2 (residues 36-46), *α*B (residues 56-101), *α*C (residues 106-176), *α*F (residues 208-259) and *α*G (residues 269-291). The segment with residues 177-203 is referred to as the *β*-tongue. The residues in *β*-tongue are flexible and highly hydrophobic in nature. The overall length of the protein is ∼10 nm forming a 34 kDa protein. An energy minimized atomistic configuration of the ClyA monomer was obtained from the work of Sathyanarayana *et al*., ^96^ where homology modelling was used to determine the structure of the N and C termini segments. The equivalent CG-mapped structure was simulated in water using the DPD force field. The residue-residue contact maps within a cutoff distance are depicted in Figs. 10A and B for the DPD as well as AA models of ClyA. The DPD model is able to faithfully reproduce most of the contacts observed in the AA data. We also observed additional contacts formed between residues 50-100 (*α*B) and 200-250 (*α*F) in the DPD simulations (Figs. 10A) which form part of the helical bundle in the ClyA monomer.

**Fig. 10.**
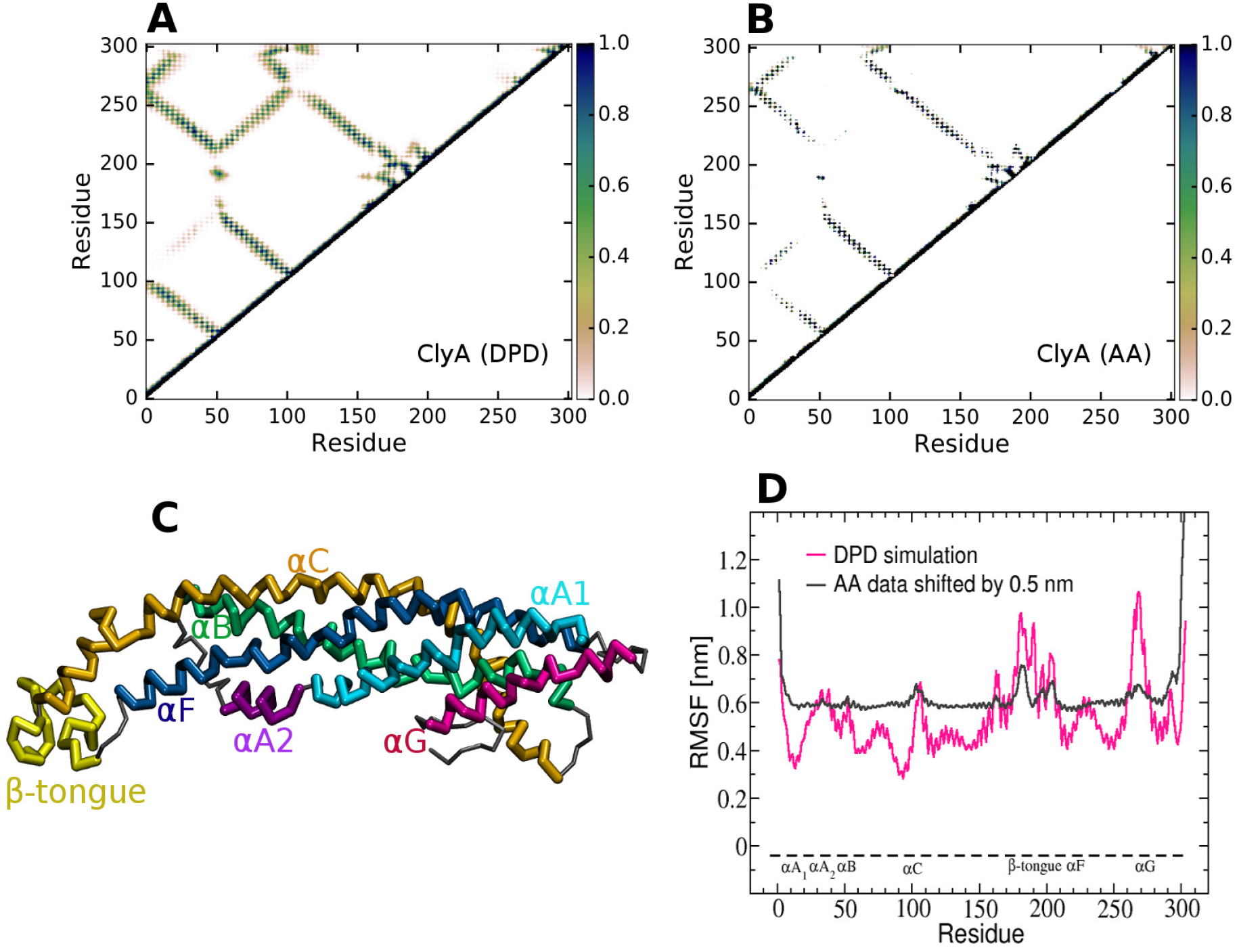
The performance of DPD force field for cytolysin A (ClyA) monomer in water and its comparison with AA simulation results. (A) The residue-residue contact map with a cutoff distance of 1.2 nm between pairs of backbone beads in the DPD simulation. (B) Contact maps constructed from AA simulations for the pairs of C_*α*_ atoms with a distance cutoff of 1.0 nm. (C) The final snapshot indicates the backbone of ClyA, with the color scheme similar to that used in Fig. 1 for ClyA. (D) Root mean square fluctuations of the backbone bead positions for ClyA from both the DPD and AA simulations.

For most of the ClyA residues, the RMSD (Fig. S6 in ESI) as well as the RMSF (Fig. 10D) are within 0.5 nm, which is lower than the length scale of the CG beads. In AA model, an increased RMSF is observed in the loop regions that connect the different helices. In addition a large RMSF is observed in the *β*-tongue residues where a large conformational changes occur upon binding with a phospholipid membrane. Interestingly our DPD model is able to qualitatively capture the trends in RMSF observed in the AA simulation for ClyA, however the base line fluctuations are larger in the DPD model. Fig. 10C illustrates that the secondary structures are preserved in the DPD simulation of ClyA with an overall compactness similar to that observed in the crystal structure (Fig. 1). Overall our results indicate that the proposed DPD parametrization is able to capture and reliably retain the secondary structure of ClyA when compared with the AA model.

## 6 Conclusions

We propose a CG force field for simulating peptides and proteins within the DPD framework. The bonded interactions in the DPD force field were parametrized using representative small peptides consisting of an *α*-helix, *β*-sheet and the fusion peptide of SARS-CoV-2 having an extended loop structure. The performance of the proposed DPD force field was extensively tested and compared with AA simulations using other short peptides, a large 34 kDa protein with 303 residues found in the pore-forming protein ClyA and the lysozyme protein having 129 residues. When compared with fully atomistic simulations, the proposed DPD force field is found to reliably preserve the protein structure and compactness as measured by the RMSD and *R*_*g*_ respectively. Excellent agreement of the native contacts with AA simulations further validates the accuracy of the proposed DPD force field. Additionally, the conformations sampled on two-dimensional Ramachandran-like *θ* − *ϕ* maps were in agreement with similar maps used in a previous CG-MD study ^92^ for the respective secondary structures.

With the ability to retain the native protein structure, the model captures protein folding from unfolded states. The dihedral potential is a key factor in stabilizing secondary conformations of the different protein structures. The dual-basin potential for the protein backbone angles, which is an improvement over the previous DPD models, effectively covers the entire spectrum of secondary structures consisting of helices, loops and extended conformations.

When compared with existing DPD models for proteins, ^52–56,58^ our proposed DPD force field reliably stabilizes a variety of protein secondary structures, reproduces native contacts, and captures protein folding to the native state. Our model alleviates the limitations present in previous Morse potential based DPD models ^52–54^ that have been used to predict secondary structures of proteins, and obviates the need to reparametrize potentials for specific protein structures. Comparing our model with polarizable DPD force fields, ^55^ improved RMSD values are obtained for the Trpcage (helix-turn-loop motif) and Trpzip (*β* sheet motif) using the proposed DPD force field. Unlike an elastic network model that controls the structural flexibility by the use of harmonic potentials between the pairs of the backbone beads, ^56^ our protein model retains the backbone flexibility by incorporating backbone angle and dihedral potentials parametrized with AA reference models. We therefore expect our model to be computationally efficient, especially for simulating large proteins and membrane-protein domains. The trimeric spike protein of SARS-CoV-2 can also be now simulated using our DPD force field.

Combined with the backbone angle and dihedral potentials, our proposed DPD force field, parametrized using the Flory-Huggins parameters for 20 standard amino acids, is generic and not limited to any specific secondary structure. We finally point out that the strength of the repulsion parameters used in our study are compatible with the existing repulsion parameters used for phospholipid membranes, allowing one to study protein-membrane systems without significant reparametrization.

## Acknowledgements

We gratefully acknowledge Thematic Unit of Excellence on Computational Materials Science (TUE-CMS) for the computational facilities. One of us, Rakesh Vaiwala acknowledges support from Unilever Research and Development (Bengaluru, India). We also thank Rochish Thaokar and Soumava Palit from Indian Institute of Technology Bombay (IITB) for initial discussions on protein simulations using the DPD methodology. We also thank Avijeet Kulshrestha from our laboratory for providing the all-atom simulation data for ClyA.

