## Supplementary Information for "A Generic Force Field for Simulating Native Protein Structures Using Dissipative Particle Dynamics"

Electronic Supplementary Information

Rakesh Vaiwala,[a] K. Ganapathy Ayappa [a,b]

[a] *Department of Chemical Engineering, Indian Institute of Science, Bangalore 560012, India*
[b] *Centre for BioSystems Science and Engineering, Indian Institute of Science, Bangalore 560012, India*

### List of Figures

S1. Probability histograms for root mean square deviation (RMSD) and radius of gyration ($R_g$) for peptides 1HDN, 1J4M and SARS-CoV-2 (6VSB).

S2. Residue-residue contact maps for the peptides 1HDN, 1J4M and 6VSB.

S3. The nativeness order parameters for the peptides 1L4X, Trpzip and Trpcage.

S4. Ramachandran-like plots on the scales of backbone angles $\theta$ and $\phi$ for the peptides 1HDN, 1J4M and 6VSB.

S5. The root mean square deviations (RMSD) and root mean square fluctuations (RMSF) for the lysozyme protein.

S6. The root mean square deviation (RMSD) for cytolysin A (ClyA) monomer.

### List of Tables

S1. Dihedral angles $\phi_0$ in the native structures, computed using the positions of four consecutive $C_\alpha$ atoms along peptide backbones.

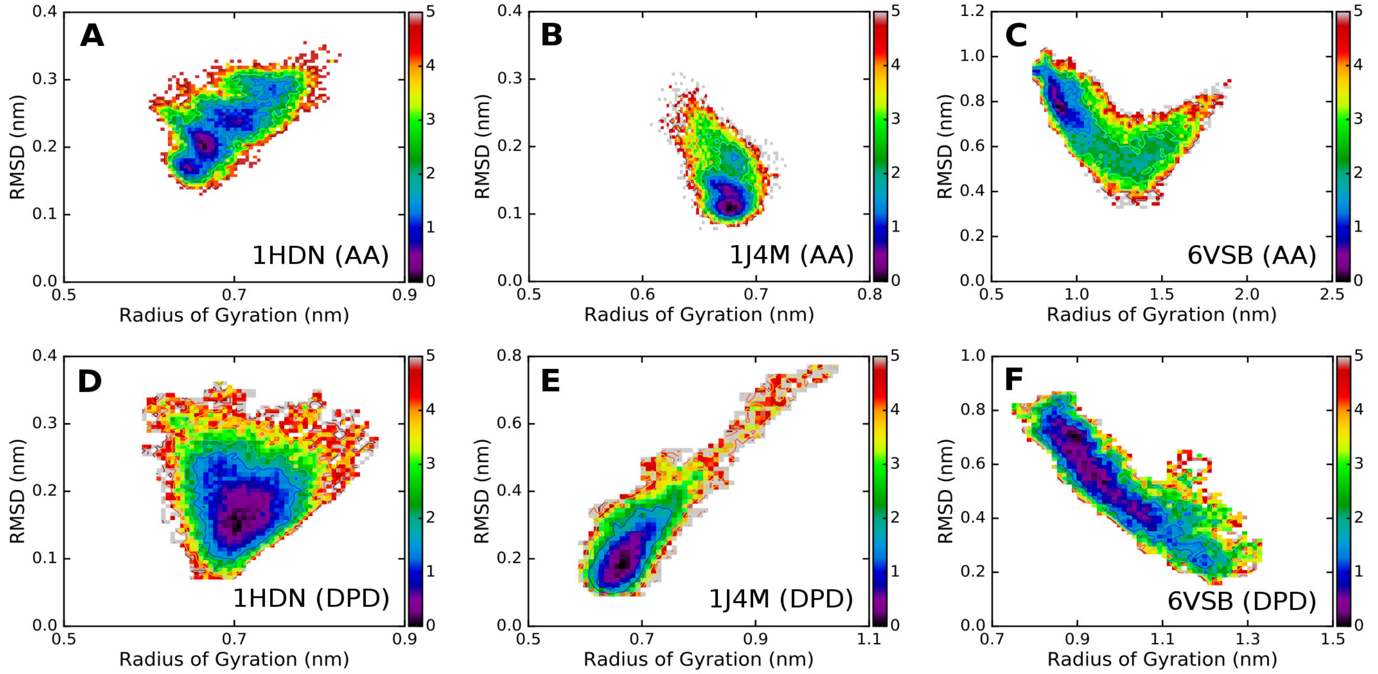


**Figure S1:** Color maps indicate the joint probability histograms $P$ for the root mean square deviation (RMSD) and radius of gyration on the log-normal scale of $-\log(P/P_{max})$, where $P_{max}$ is the maximum value of $P$ in histograms. The histograms (A, B, C) refer to the all-atom (AA) simulations for peptides 1HDN (A), 1J4M (B) and 6VSB (C), while the corresponding DPD results are shown by the panels (D), (E), and (F), respectively.

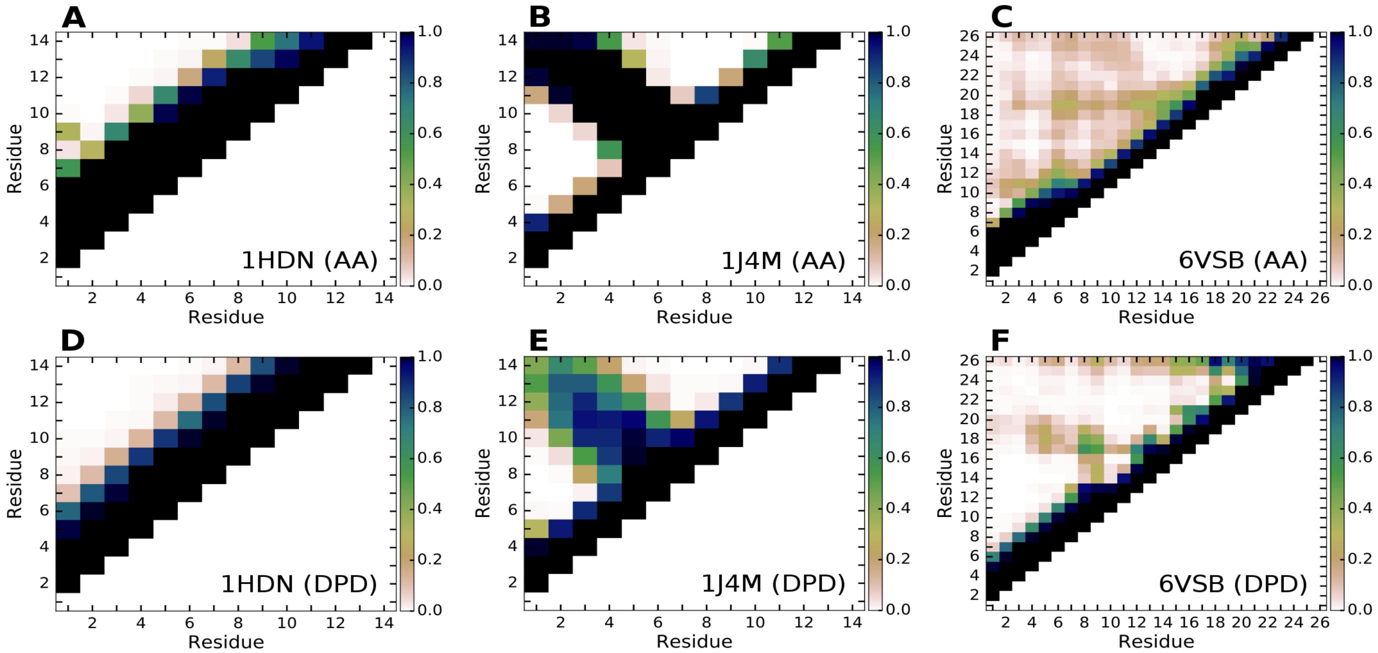


**Figure S2:** The contact maps with a cutoff distance of 1.0 nm are shown for the peptides 1HDN (A, D), 1J4M (B, E) and 6VSB (C, F). The color bar indicates a fraction of the trajectory wherein the contacts are stabilized. The maps obtained from the AA simulations are indicated in panels (A), (B) and (C), while those derived from the DPD simulations are illustrated in panels (D), (E) and (F).

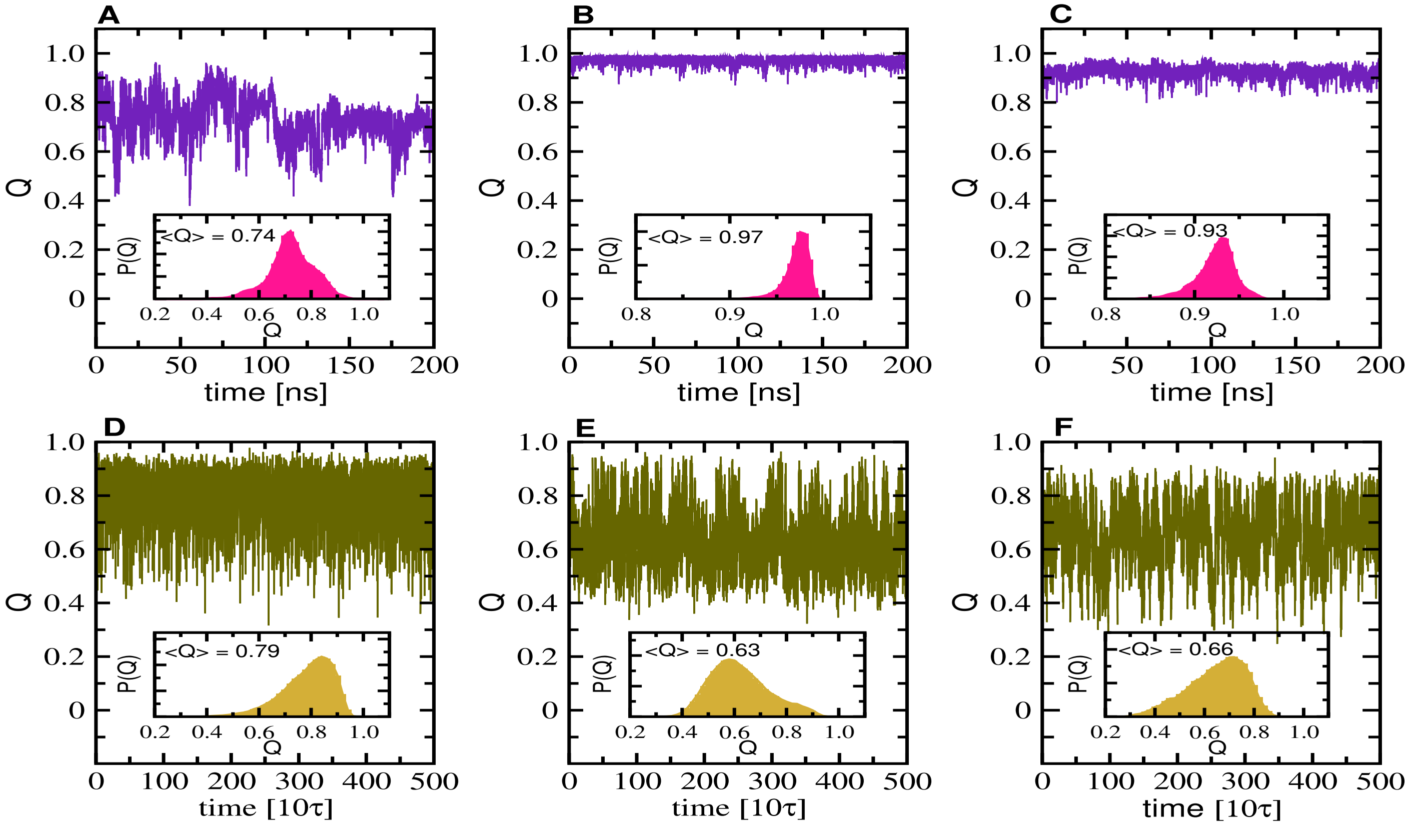


**Figure S3:** The temporal variation of the order parameter $Q$ (see main text for definition) for the model peptides 1L4X (A, D), Trpzip (B, E) and Trpcage (C, F). The top panels (A, B, C) are obtained from AA simulations, while the panels at the bottom (D, E, F) are derived from the DPD simulations. The insets show the histograms for the order parameter, and $\tau$ is the DPD time scale.

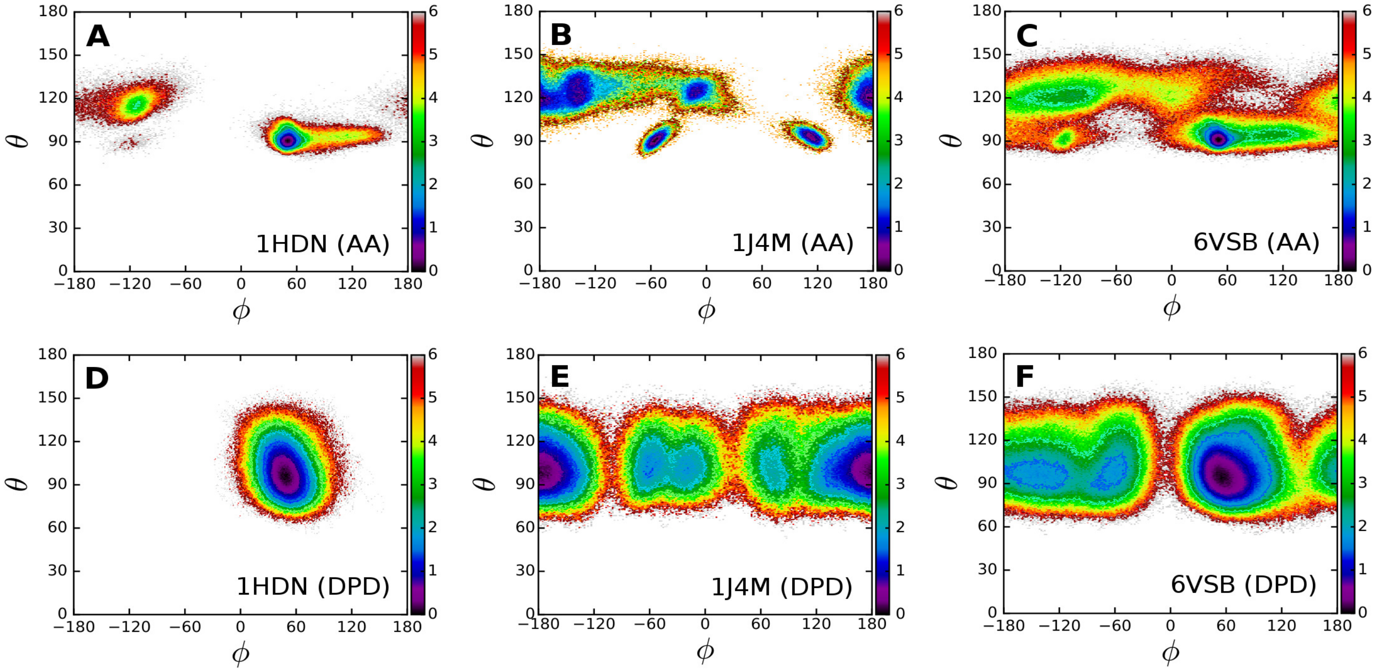


**Figure S4:** Two dimensional plots for the model peptides 1HDN (A, D), 1J4M (B, E) and 6VSB (C, F), for the probability density $P(\phi, \theta) = -\log(P(\phi, \theta)/P_{max})$ on the phase space of backbone angle $\theta$ and dihedral angle $\phi$. The maximum density is $P_{max}$. The top panels (A, B, C) are obtained from AA simulations, while the panels at the bottom (D, E, F) are derived from the DPD simulations. The angles are represented in degrees.

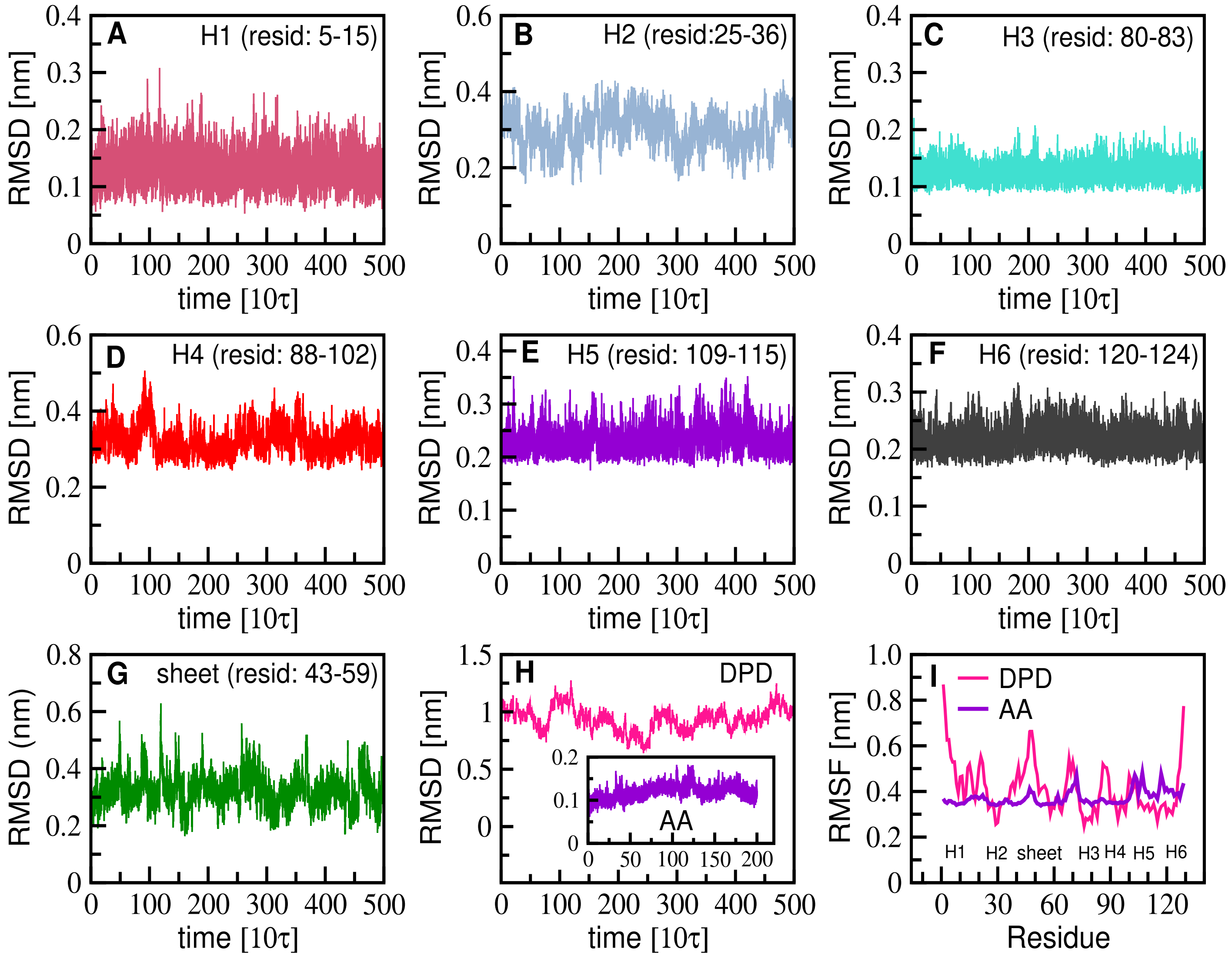


**Figure S5:** (A-G) Root mean square deviation (RMSD) for the structural segments of lysozyme (PDB:3TXJ) simulated in water using the proposed DPD force fields. The RMSDs of six helices (H1-H6) and segment with extended sheet are stabilized at $\sim 0.3$ nm in the DPD simulation. (H) The overall RMSD for lysozyme is $\sim 0.94$ nm in DPD simulation due to flexible loops and turns in the protein, while it is $\sim 0.1$ nm in all-atom (AA) simulation indicated in the inset. (I) The root mean square fluctuations (RMSF) of amino acid residues are within to $\sim 0.5$ nm for most of the protein. The RMSF for the lysozyme simulated with AA force fields is shifted by 0.3 nm for ease of comparison. The symbol $\tau$ denotes the DPD time scale. The last half trajectory is used for analysis.

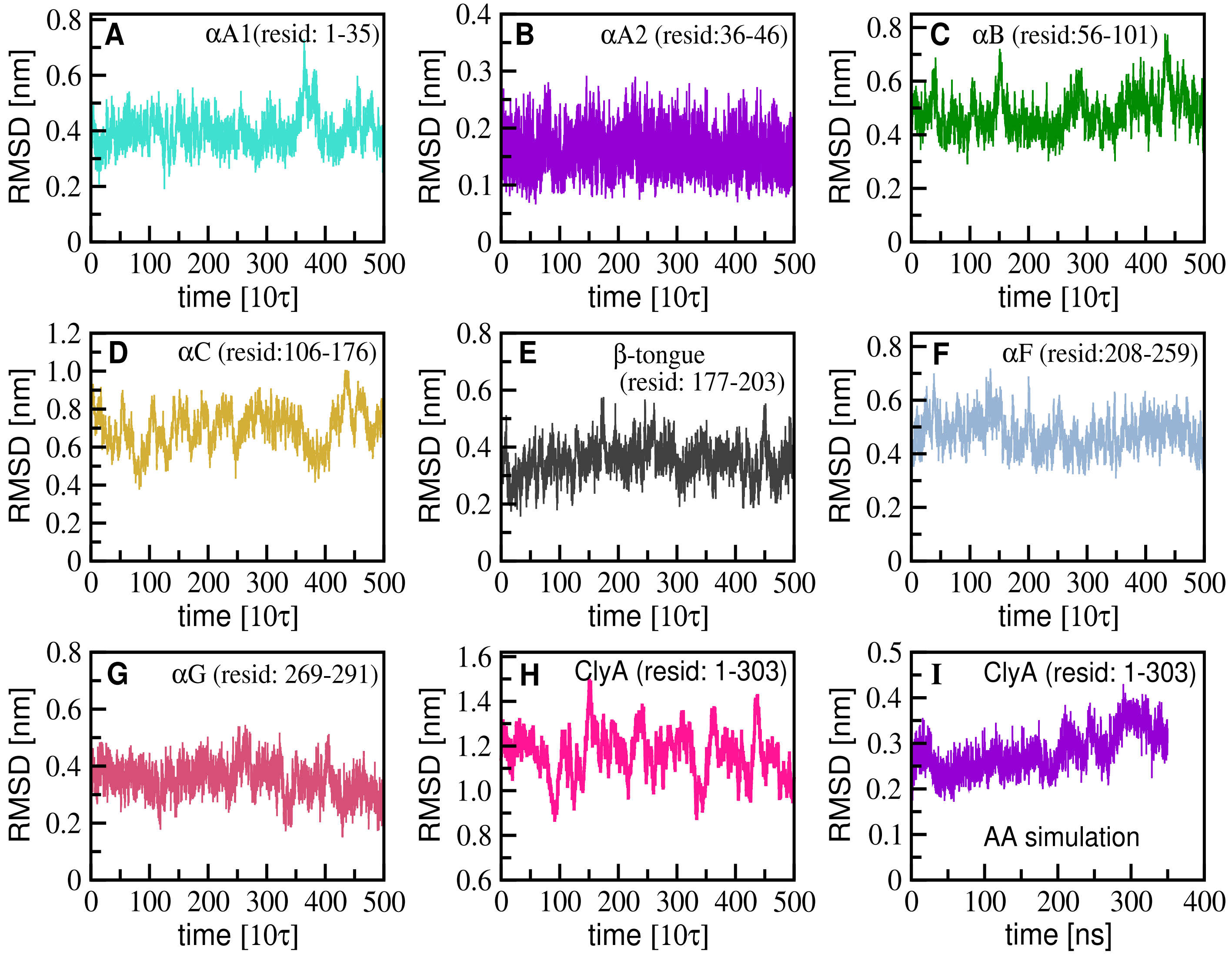


**Figure S6:** Root mean square deviation (RMSD) for the different structural segments of the ClyA monomer simulated in water using DPD simulations (A-H) and AA simulations (I). Most of the segments show a RMSD below 0.5 nm (A-C and E-G) while the segment $\alpha$C is relatively more flexible with RMSD $\sim 0.7$ nm in DPD simulation (D). The RMSD for the entire ClyA protein is stabilized at $\sim 0.3$ nm in AA simulation (I), while it is quite flexible in DPD simulation due to $\alpha$C helix and flexible loops (H). The symbol $\tau$ denotes the DPD time scale. The DPD simulation trajectory comprises of $10^6$ steps with $dt = 0.01\tau$, and the temporal variation for the last half of the trajectory is indicated.

**Table S1:** Dihedral angles $\phi_0$ (degree unit) in the different native structures, computed using the positions of four consecutive $C_\alpha$ atoms $i-l$ along the peptide backbones.

| Dihedrals ($C_\alpha$ atoms $i-l$) | Peptides (residues) | | | | | |
|---|---|---|---|---|---|---|
| | **1HDN** (14) | **1J4M** (14) | **6VSB** (26) | **1L4X** (15) | **Trpzip** (12) | **Trpcage** (20) |
| $1-4$ | 56.16 | 76.24 | 48.95 | 47.00 | 177.98 | 45.80 |
| $2-5$ | 46.43 | 173.41 | 53.27 | 48.29 | -108.68 | 55.26 |
| $3-6$ | 43.95 | -171.91 | 53.46 | 51.38 | 170.41 | 49.32 |
| $4-7$ | 57.42 | 166.00 | 49.17 | 45.40 | -36.58 | 48.41 |
| $5-8$ | 42.01 | -1.68 | 47.23 | 49.50 | -29.83 | 46.80 |
| $6-9$ | 48.55 | -59.33 | 53.86 | 52.10 | 70.08 | 59.69 |
| $7-10$ | 51.35 | 127.68 | 44.53 | 50.10 | -114.35 | 56.69 |
| $8-11$ | 54.14 | -148.05 | 72.90 | 47.21 | -168.96 | -87.29 |
| $9-12$ | 41.42 | -175.57 | 53.16 | 53.62 | -113.29 | -127.53 |
| $10-13$ | 54.28 | -165.73 | -136.69 | 54.55 | | 4.41 |
| $11-14$ | 44.10 | 176.67 | -140.13 | 53.93 | | 66.02 |
| $12-15$ | | | -115.73 | 45.65 | | 70.33 |
| $13-16$ | | | -76.08 | | | -109.02 |
| $14-17$ | | | 97.46 | | | 103.82 |
| $15-18$ | | | 101.15 | | | -122.33 |
| $16-19$ | | | 82.02 | | | -90.51 |
| $17-20$ | | | -163.12 | | | -108.42 |
| $18-21$ | | | 68.73 | | | |
| $19-22$ | | | -174.16 | | | |
| $20-23$ | | | 79.03 | | | |
| $21-24$ | | | 71.61 | | | |
| $22-25$ | | | -54.52 | | | |
| $23-26$ | | | -48.16 | | | |
